# ProteomeScan: A Toolkit For Target Validation By Proteome-Wide Docking And Analysis

**DOI:** 10.64898/2026.04.14.718479

**Authors:** Aryan Amit Barsainyan, Rishikesh Panda, Jose Siguenza, Daniele Merico, Bharath Ramsundar

## Abstract

The problem of identifying which protein target a potential drug-like molecule interacts with is crucial for both the study of existing drugs and the design of new therapeutic compounds. Despite the importance of target identification, existing computational approaches remain limited in terms of speed, accuracy, and protein target coverage. We introduce ProteomeScan, a large-scale, gene-driven computational toolkit for systematic proteome-wide scanning to uncover hidden or previously uncharacterized protein-ligand interactions. ProteomeScan leverages cloud-scale high performance computing to perform extensive molecular docking simulations across the human proteome to rank candidate targets based on binding affinities. After filtering promiscuous targets, we found that ProteomeScan ranks known target significantly better than a random baseline for a set of control compounds. Furthermore, we performed physical analyses of predicted binding modes for both promiscuous and known protein-ligand binding pairs to validate that ProteomeScan identifies interactions with valid binding pockets. In addition, we conducted experiments using mutant variants of proteins to study how mutations affect binding behavior. We have open sourced the core ProteomeScan algorithm as part of the DeepChem ecosystem to enhance transparency and reproducibility.

**Author summary:** 

## Introduction

Target identification is a critical step in drug discovery and the study of disease mechanisms as it bridges the gap between chemical compounds and their biological effects [1]. By simulating the interactions between molecular targets, such as proteins or enzymes, and a drug candidate, researchers gain insight into the compound ’s mechanisms of action, therapeutic potential, and possible side effects [2]. Also, target identification enables drug re-purposing, where existing compounds are evaluated for new therapeutic applications, significantly reducing development costs and time [3].

Current methods for computational target identification often focus on a limited subset of targets, potentially leading to missed therapeutic opportunities or the failure to uncover off-target effects [4].

We aim to systematically scan the human proteome for drug-protein interactions. We have developed ProteomeScan, a computational toolkit designed for large-scale, full-proteome scanning that starts with selection optimal PDB entries for each gene in the human proteome combined with physics-based docking algorithms [5]. We conduct a comprehensive search of the human proteome using the UniProt reference database [6] using the data-preparation pipeline. After filtering for reviewed proteins with verified protein-level existence and removing duplicates and irrelevant entries, we finalize a dataset comprising 7,657 unique genes. All the gene-products are assessed for potential interactions against 20 selected ligands for each selected gene. Table 1 shows the list of 20 ligands selected for the run and their known targets that were used for positive control and downstream analysis. We mainly selected signaling inhibitors, primarily targeting the RTK/MAPK/AKT pathways (e.g. alpelisib, dabrafenib, erlotinib, trametinib) but also a few other targets/pathways (e.g. palbociclib, navitoclax), plus a few DNA- and microtubule-targeted chemotherapeutic agents (olaparib, paclitaxel, SN-38). We preferentially selected approved drugs, but also included investigational ones with a well-characterized mechanism of action. Finally, we ensured the presence of non-kinase targets.

**Table 1.** The table lists the 20 ligands used for the ProteomeScan run and their known targets were used for positive control and downstream analysis. We mainly selected signaling inhibitors, primarily targeting the RTK/MAPK/AKT pathways (e.g. alpelisib, dabrafenib, erlotinib, trametinib) but also a few other targets/pathways (e.g. palbociclib, navitoclax), plus a few DNA- and microtubule-targeted chemotherapeutic agents (olaparib, paclitaxel, SN-38). We preferentially selected approved drugs, but also included investigational ones with a well-characterized mechanism of action. Finally, we ensured the presence of non-kinase targets.

| Compounds | Target_symbol |
| --- | --- |
| Alpelisib | PIK3CA |
| Binimetinib | MAP2K1, MAP2K2 |
| Dabrafenib | BRAF, RAF1 |
| Erlotinib | EGFR |
| Navitoclax | BCL2, BCL2L1, BCL2L2 |
| Olaparib | PARP1, PARP2 |
| Paclitaxel | TUBB |
| Palbociclib | CDK4, CDK6 |
| Regorafenib | RAF1, KIT, PDGFRB, RET, FLT1, KDR, FLT4 |
| RMC-6236 | KRAS, NRAS, PPIA, HRAS |
| Sapanisertib | MTOR |
| Selinexor | XPO1 |
| SN-38 | TOP1 |
| SOS1-IN-11 | SOS1 |
| SOS1-IN-15 | SOS1 |
| TAK-632 | BRAF, RAF1 |
| Trametinib | MAP2K1, MAP2K2 |
| Tucatinib | ERBB2 |
| Vemurafenib | BRAF, RAF1 |
| Vociprotafib | PTPN11 |

We find that large-scale scanning across the proteome can be complicated by an effect we call protein promiscuity, wherein a single protein interacts with multiple ligands. Off-target interactions from protein promiscuity can obscure true targets and may possibly lead to unwanted side effects in drug discovery [7]. Because a promiscuous protein may rank highly for many compounds regardless of true specificity, failing to account for these broad binders can obscure genuine high-affinity targets and lead to false positives in large-scale screens. To address this, we conducted a systematic promiscuity analysis by identifying proteins that repeatedly appear among top docking hits for different ligands, allowing us to quantify how shared off-target interactions affect our scoring and filtering. By separating broadly binding proteins from those with specific high-affinity interactions, we improve the precision of target validation within ProteomeScan’s proteome-wide framework.

The ProteomeScan target validation toolkit has been open-sourced through the DeepChem server, part of the broader DeepChem ecosystem [8], an open-source library that provides datasets, models, and tools for applying machine learning to chemistry, biology, and materials science. DeepChem has introduced widely used tools including the MoleculeNet [9] benchmark suite and the ChemBERTa [10] series of chemical foundation models. A scaled version of ProteomeScan is also available in the commercially available Prithvi software suite. By making proteome scanning software widely accessible, we aim to accelerate the discovery of new biology and support the development of innovative therapeutics.

### 1 Related Work

Computational target identification has seen remarkable advances driven by breakthroughs in structural biology, machine learning, and high-performance computing. A broad range of computational target-identification algorithms have been proposed in past literature. We briefly survey these methods here. We discuss these past efforts, successes and their limitations below.

#### 1.1 Target Identification Methods

Computational target identification approaches can be categorized into two primary categories: structure-based methods that use three-dimensional protein structures to predict drug-target interactions, and ligand-based methods that rely on chemical similarity principles using known bioactive molecules.

Traditional molecular docking follows a “many compounds-one target” approach, predicting ligand-receptor complex structures by sampling ligand conformations in protein active sites and ranking them via scoring functions. Building upon this foundation, reverse docking introduced a paradigm shift from the traditional “many compounds-one target” virtual screening to a “one compound-many targets” approach. This approach enabled systematic exploration of drug polypharmacology by screening individual compounds against comprehensive protein databases rather than focusing on single targets. Reverse docking emerged with tools like TarFisDock [11], a structure-based platform that established a proof-of-concept by using a database of 698 protein structures and demonstrated the feasibility of multi-target screening approaches, and PharmMapper [12], which used structure-based pharmacophore models extracted from 3D protein-ligand complexes for reverse mapping. However, these early platforms were constrained by limited protein coverage and lacked validation methods.

In parallel with structure-based reverse docking developments, past work has explored ligand-based approaches that improved efficiency by leveraging chemical similarity. PASS [13] introduced biological activity spectra from structural formulae, while SEA [14] related proteins through ligand similarity. ChemMapper [15] was developed as an online platform using 3D similarity computation for polypharmacology prediction, while ChemProt [16] integrated chemical-protein databases with similarity-based target prediction. SuperPred [17] refined these approaches by integrating 2D/3D similarity, and physicochemical properties with BLAST-based statistical analysis, and SwissTargetPrediction [18] achieved scalability, processing 376,342 compounds against 3,068 targets in 15-20 seconds. However, these methods cannot identify specific binding sites or validate whether predicted interactions occur in druggable pockets and are more approximate methods, as they don’t directly score a ligand based on the protein structure. Some work has explored combining ligand-based and structure-based approaches: LigTMap [19] demonstrated the effectiveness of integrating similarity searches with molecular docking for target prediction.

Researchers pursued scaling up structure-based approaches along a path toward proteome-wide coverage, driven by the need to preserve critical binding site information. ReverseDock [20] enabled blind docking of single ligands against up to 100 user-selected proteins, ACID [21] implemented consensus inverse docking combining multiple scoring functions for improved accuracy, and CRDS [22] performs reverse docking against over 5,000 protein structures using consensus scoring. While consensus approaches enhance hit identification, they significantly increase computational costs by requiring simultaneous execution of multiple docking programs. At the proteome scale, these computational demands become prohibitive, as structure-based methods like molecular docking are already time intensive. Lee and Kim’s [23] pioneering study performed reverse docking against 10,886 human binding sites, establishing that comprehensive protein structure coverage is crucial for identifying potential targets. More recently, during the Covid-19 pandemic, researchers identified 22 FDA-approved drug candidates by using reverse docking against SARS-CoV-2 targets [24]. Kong et al. [25] screened 33,446 predicted binding pockets across 80% of the AlphaFold human proteome, successfully identifying folate receptors as key PFAS targets in the first proteome-wide reverse docking study in environmental toxicology.

These studies establish the scientific foundation and practical feasibility of proteome-wide reverse docking. ProteomeScan approaches proteome-wide reverse docking through systematic gene-based structure selection, comprehensive protein promiscuity analysis, and druggable pocket validation in a unified pipeline. In addition, ProteomeScan introduces a proteome-coverage–level evaluation metric, termed *Known Target Recovery (KTR)*, to quantify how effectively deep docking functions can recapitulate known target–ligand interactions at scale.

#### 1.2 Protein Promiscuity

One of the key findings using ProteomeScan in this work is the identification of a class of proteins which appear to bind a broad range of different ligands. This phenomenon, known as protein promiscuity, is central to computational drug discovery, presenting both a critical challenge and a significant opportunity [26]. We briefly survey past work on protein promiscuity to provide a scientific base for subsequent discussion.

Proteins bind to multiple ligands for several key reasons. Conformational flexibility allows a binding site to adopt different shapes [7]. Allosteric effects propagate changes from distant sites into the active site, broadening ligand scope [27].

Partial recognition allows imperfect fits to still bind, enabling activity on related substrates [28]. Multiple interaction sites let one protein accommodate diverse ligands either in the same pocket (different orientations) or at distinct sites, a behaviour common in antibodies [29]. These mechanistic insights explain why some proteins consistently appeared as top hits in large-scale docking studies, regardless of the specific compound being screened. Understanding this pattern was crucial for distinguishing genuine high-affinity interactions from artifacts of promiscuous binding.

Large-scale structural analysis indicates that more than one-third of representative binding pockets from the Protein Data Bank are promiscuous and interact with multiple, chemically different ligands [30]. Protein promiscuity is far more common than previously recognized and has important implications for biotechnology and drug discovery. Ignoring protein promiscuity effects in proteome-scale docking leads to an overwhelming number of false positives and confounds target prioritization, as broadly binding proteins obscure specific interactions. ProteomeScan addresses this by making promiscuity analysis central to the target identification process, enabling both improved specificity and insights into polypharmacological opportunities and off target identification.

## 2 Methods

ProteomeScan, illustrated in figure 1(A), automates proteome-wide target validation by first assembling a curated library of canonical human protein structures and then running parallelized AutoDock Vina simulations against over 7,500 gene products.

**Fig 1.**
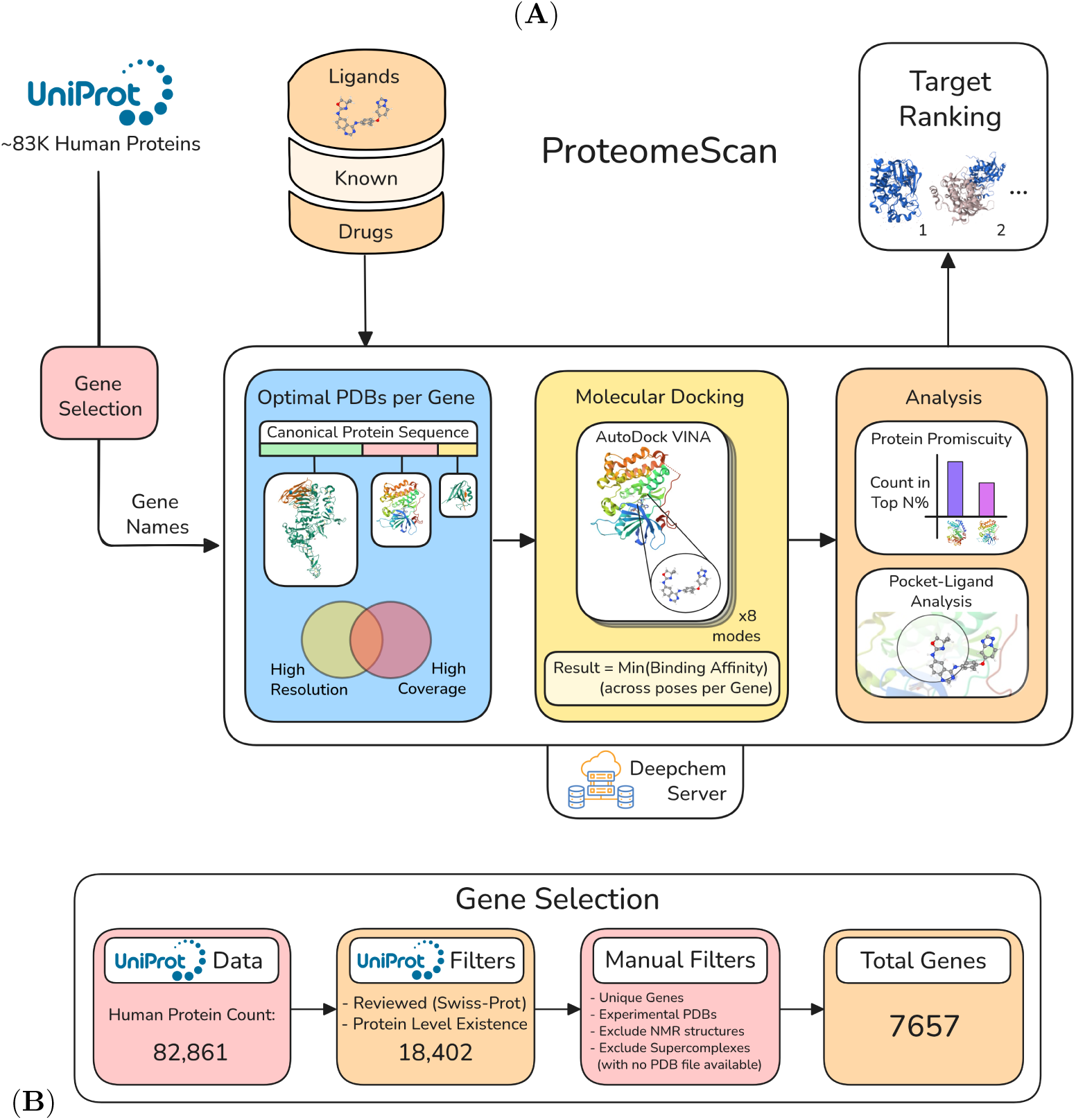
(A) ProteomeScan Overview: The diagram illustrates the workflow for the ProteomeScan toolkit available on the Deepchem Server. The pipeline takes as input human proteome gene names and known drug ligands. In the first stage, the pipeline selects optimal PDB structures corresponding to each gene’s canonical protein sequence. These structures are chosen to have high resolution (*<* 2.5–3 °A) and to collectively provide maximum sequence coverage. The second stage performs molecular docking for all ligand-PDB structure combinations associated with each gene. For each gene, the lowest binding affinity score across all its PDB structures is retained. This step is conducted across the entire proteome. In the third stage, the docking results are examined. The top-ranked genes for each ligand are compared against rankings from known drug-target pairs to identify potential promiscuous targets. Furthermore, both the top-ranked promiscuous and non-promiscuous genes undergo analysis to evaluate the extent of druggable pocket coverage by the ligands. After applying specific filtering criteria and re-ranking, the ProteomeScan narrows down the top targets. (B) Data preparation for full human proteome scan: We retrieved 82,861 human protein entries from UniProt, which provides reviewed sequences and gene annotations. To ensure one representative sequence per gene, we collapsed multiple PDB structures to a single canonical protein per gene. We excluded entries without experimental PDB data, including NMR models, and filtered out structures with exceptionally large atom counts. This resulted in 7,657 unique genes with canonical structures for proteome-wide docking analysis.

Docking scores are aggregated and subjected to a systematic promiscuity analysis, which filters out broadly binding proteins and highlights true high-affinity targets. The entire workflow, from structure selection through score processing, is exposed via the RESTful API on the open-source DeepChem server, with scalable execution managed by the commercial Prithvi suite. For the detailed algorithm, please refer to 1.

### 2.1 Data Preparation

We utilized the UniProt database to obtain 82,861 human protein entries. UniProt provides thoroughly reviewed sequences with comprehensive annotations, including amino acid sequences and corresponding gene information for human proteins. Since a single gene may be associated with multiple PDB structures, we consolidated the data by selecting one canonical protein sequence per gene to ensure a consistent and representative dataset. Next, we excluded entries without experimental PDB structures, including all NMR-derived models, and additionally excluded PDB IDs for supercomplexes with extremely large atom counts, as PDB format files were unavailable. Following these curation steps, we retained 7,657 unique genes, along with their canonical protein structures, which formed the foundation for our systematic, proteome-wide docking analysis. This step is illustrated in figure 1(B) and a detailed description of the data preparation algorithm 2 can be found in Appendix.

#### Protein Retrieval and Sequence Data

For each of the 7,657 genes, a list of PDBs was obtained from data sources such as PDBe-KB [31]. Each PDB entry was associated with coverage and resolution data. The PDB files represented varying fragments of protein, and we assessed their quality based on the resolution and the extent of sequence coverage.

#### Selection of Optimal PDB Entries

A selection algorithm is applied to choose the most optimal PDB entries for each gene. The criteria for selection include the resolution and sequence coverage of each structure. The output is a list of selected PDB entries with optimized coverage and resolution. A detailed description of the selection algorithm 3 can be found in Appendix.

#### Cleaning the Selected PDBs

After selecting the optimal PDBs, the structures undergo cleaning to remove water molecules, heteroatoms, and side chains that are unrelated to the gene of interest. Hydrogens are added to the cleaned PDBs to standardize the structures at a specific pH (typically pH 7) for docking simulations. A detailed description of the PDB cleaning algorithm 4 can be found in Appendix.

#### Preparation of Ligands

Ligand structures were generated from their corresponding SMILES representations using RDKit package. Hydrogen atoms were added to ensure proper valency and protonation states. Three-dimensional conformations were generated using the ETKDG (Experimental-Torsion Knowledge Distance Geometry) algorithm, which combines distance geometry with knowledge of experimental torsional preferences. The resulting 3D, protonated ligand structures were saved in SDF format for downstream analysis.

Given the large number of experimental structures per target, we used blind docking to avoid the need for per-structure pocket identification. Predicting and selecting binding sites across multiple PDBs per gene poses significant challenges in consistency at this scale. Blind docking enabled automated, uniform sampling of the protein surface without manual intervention, making it more practical for our pipeline. Importantly, experimentally determined structures were prioritized as they provide a more grounded and biologically validated baseline for docking performance.

Later on, we also evaluated AlphaFold-predicted structures for proteome-wide docking, however, due to their large size, multi-domain architecture, and variable pLDDT scores, they were not feasible for our blind docking based approach as their full-length nature resulted in prohibitively large search spaces. Details of the preliminary AlphaFold-based structure docking analysis are provided in the Appendix.

### 2.2 Gene Guided Protein-Ligand Docking Workflow

To systematically predict potential drug-target interactions, starting at the gene level, we implemented a high-throughput docking workflow using AutoDock Vina.

For each protein–drug pair, we compiled the binding affinity scores (predicted by Vina in kcal/mol) from all docking runs against each selected PDB structure associated with that gene. We selected the lowest binding affinity score per pair, representing the strongest predicted interaction. This assumption is based on the rationale that the most favorable (i.e., lowest energy) docking score is more likely to correspond to a biologically meaningful interaction, provided the docking poses are sterically and chemically plausible.

The final output was an aggregated dataset in CSV format, containing the best docking score for every gene–drug pair. These results serve as a foundation for downstream analyses, including candidate prioritization, promiscuity, and potential reposition insights. A detailed description of the gene-guided protein-ligand docking algorithm 5 can be found in Appendix.

### 2.3 Scalable Workflows on DeepChem Server

To accelerate the workflow, docking jobs for each gene-drug docking were run in parallel via a distributed computing infrastructure, Prithvi, built around the DeepChem Server. DeepChem Server as highlighted in fig 2, is an open-source framework for running scientific discovery pipelines at scale. We implemented Deepchem-server using FastAPI with endpoints for data handling, and workflows for machine learning, drug discovery, etc. The API supports asynchronous execution, so multiple requests can run concurrently and efficiently. Users interact with the same API to submit jobs, check status, and retrieve results.

**Fig 2.**
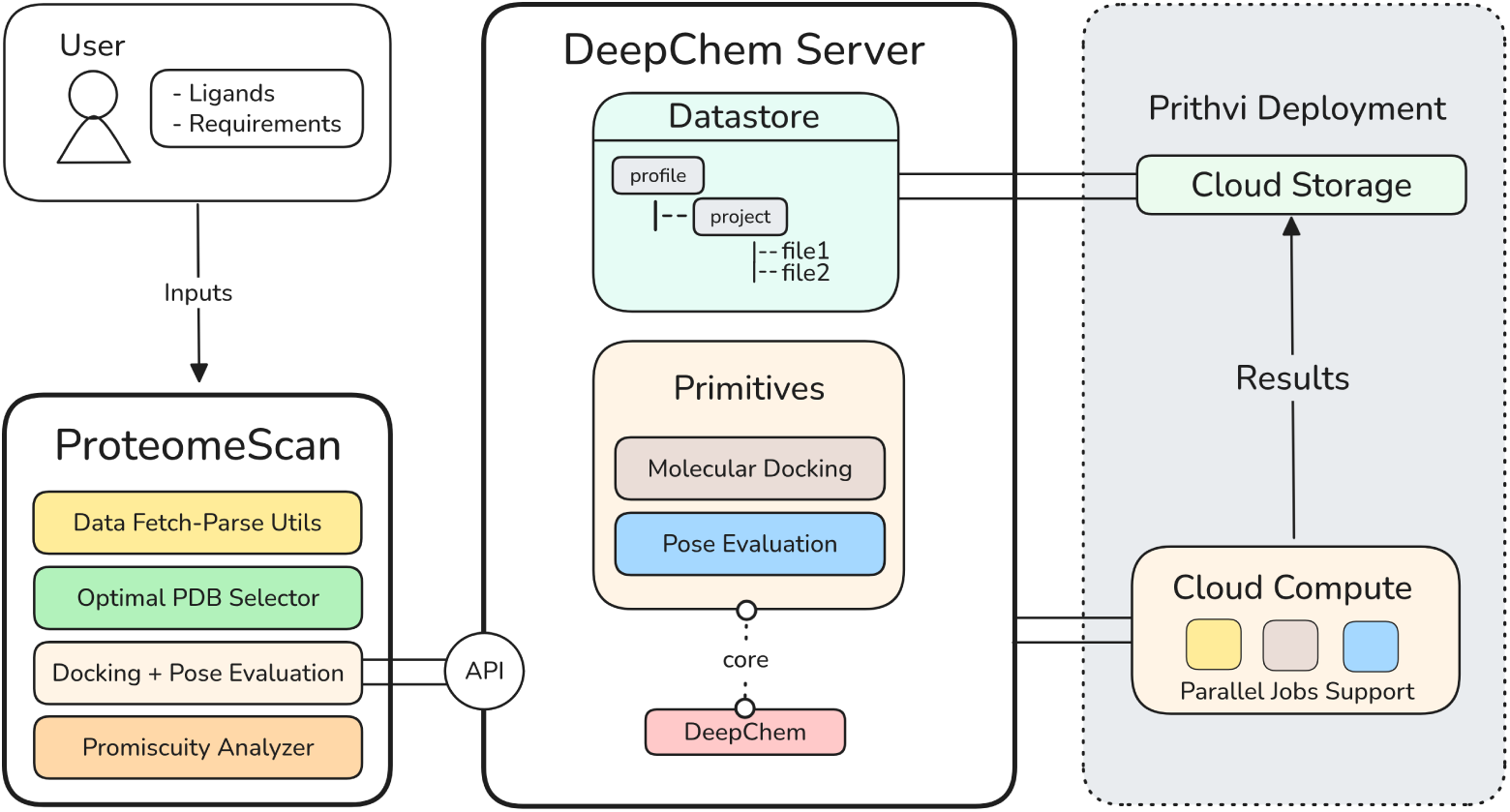
Overview of the DeepChem Server usage in ProteomeScan. Users submit data to ProteomeScan which sends workflow parameters via API, which in turn directs these requests to various DeepChem-based primitives (for example, Molecular Docking, Pose Evaluation etc.) to run the ProteomeScan. Compute-intensive jobs are dispatched to the Prithvi cloud backend for parallel execution.

Under the hood, each workflow relies on the DeepChem library to ensure reproducibility. Input files and intermediate data are stored in a local *DiskDatastore*, organized by user profile and project to keep files structured and easy to manage. A specific Deepchem address format is used to reference files within this storage system, allowing the Deepchem server endpoints to locate and use them when executing workflows. It also includes a lightweight Python SDK called *py-ds* for easy integration in Python-based clients or automation scripts. The commercial version of DeepChem server is deployed on Prithvi™ which runs DeepChem server workflows on AWS Batch.

### 2.4 Protein Promiscuity Analysis

Promiscuous targets are gene-products that exhibit high binding affinity to a majority of known drugs in the docking experiments. We observed that filtering these targets yielded better ranks for known targets in the system. Formally, promiscuous targets were identified based on the following criteria:

*A target T is promiscuous if T is in the top m*% *of targets (based on docking score) for at least n of the N ligands (with n* ≤ *N)*:

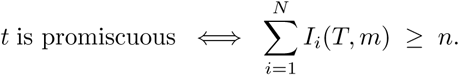

where,

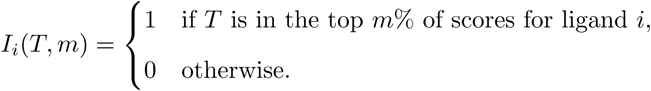

A detailed description of the promiscuity analysis algorithm 6 can be found in Appendix.

### 2.5 Pose Analysis Pipeline

We developed a pose analysis pipeline to systematically evaluate whether ligands in top-scoring complexes actually occupy druggable cavities within the protein (Figure 3A).

**Fig 3.**
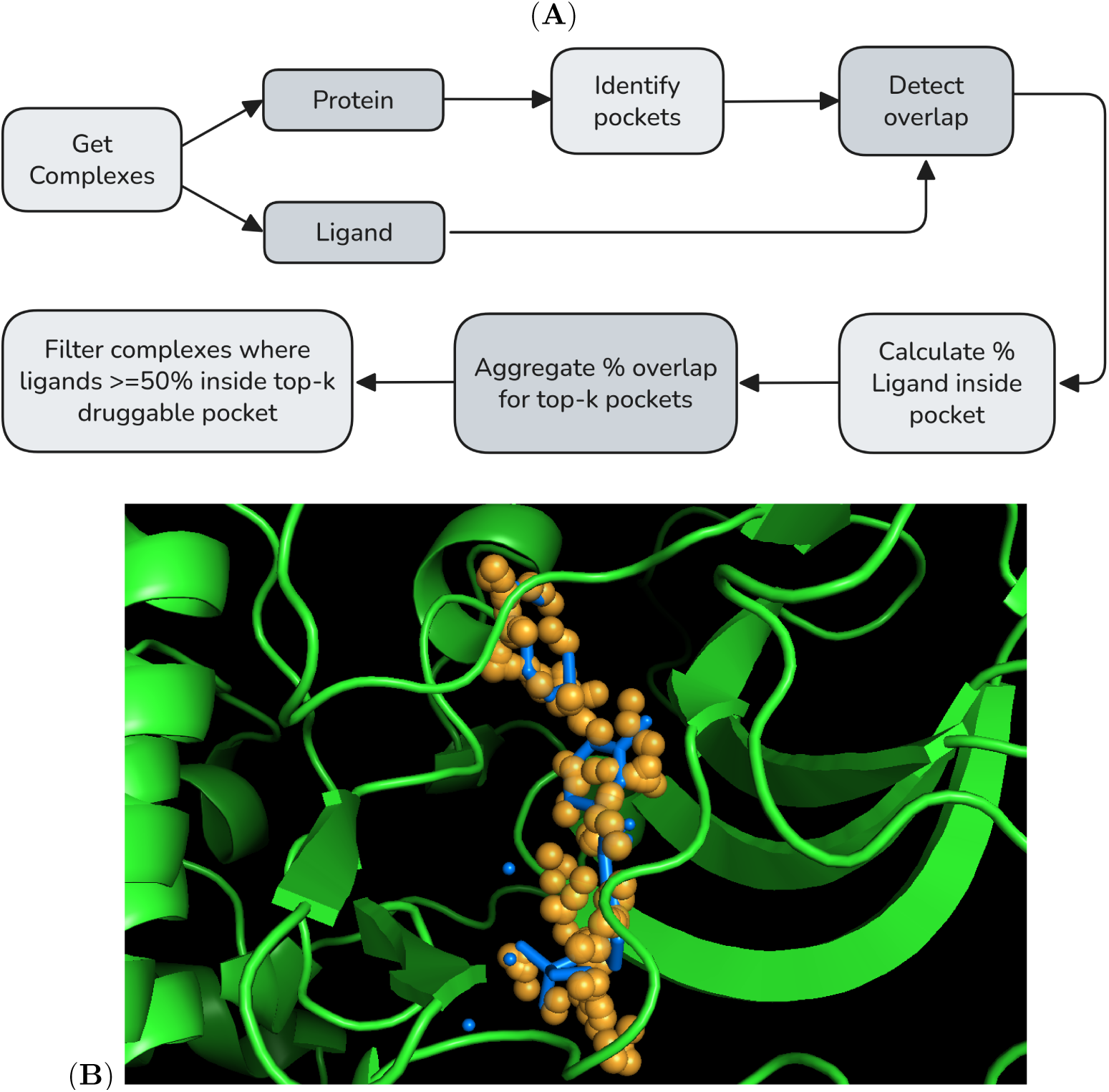
(A) Workflow for evaluating ligand-pocket overlap in protein–ligand complexes. The workflow starts with collecting complexes from docking runs. Protein structures are processed with Fpocket to identify potential binding pockets, while ligands are isolated for overlap evaluation. Overlap is quantified by computing the percentage of ligand atoms located within 3A° of Fpocket’s pocket-defining alpha-spheres. These percentages are then aggregated for the top-k pockets, ranked by Fpocket’s druggability score. Complexes are filtered to retain only those in which at least 50% of the ligand volume overlaps with a top-k predicted pocket, allowing classification of ligand binding as either in-pocket or out-of-pocket. (B) This figure shows binding pose of ERBB2 (PDB ID 8U8X, chain A) with ligand Tucatinib. Orange spheres represent fpocket detected pocket using alpha spheres and Blue docked ligand shows overlap with pocket alpha spheres. Overlap is considered when ligand atoms come within 3A distance of pocket alpha spheres.

The pipeline integrates docking results with pocket detection tools and evaluates ligand-pocket interactions based on spatial overlap and pocket quality metrics.

The analysis begins by separating each docking complex into individual protein and ligand PDB files. The protein structure is then analyzed using the fpocket tool, which identifies potential binding pockets based on geometric and physicochemical properties. Each detected pocket is represented by a set of alpha-spheres—pseudo-atomic spheres that approximate the volume and shape of cavities within the protein. Fpocket also computes key metrics such as the Druggability Score, which estimates the likelihood that a pocket is suitable for binding drug-like small molecules.

To evaluate whether the ligand resides within or overlaps with these pockets, we overlay the ligand atoms onto the pocket alpha-spheres (Figure 3B). Two custom metrics were introduced to quantify the extent and quality of ligand-pocket interactions which include “*% Pocket Filled* ” and “*% Ligand Inside Pocket* ” as described in table 2. These metrics provide a more nuanced analysis of binding poses by assessing not just placement, but the degree to which a ligand utilizes biologically relevant, druggable cavities (see example in Figure 6). We also introduced metrics like “*Total % Ligand Inside Pocket* ” and “*Total % Ligand Inside Top-n Pockets*” to facilitate cross-complex comparison, also described in table 2. A detailed description of the pose evaluation algorithm 7 can be found in Appendix.

**Fig 4.**
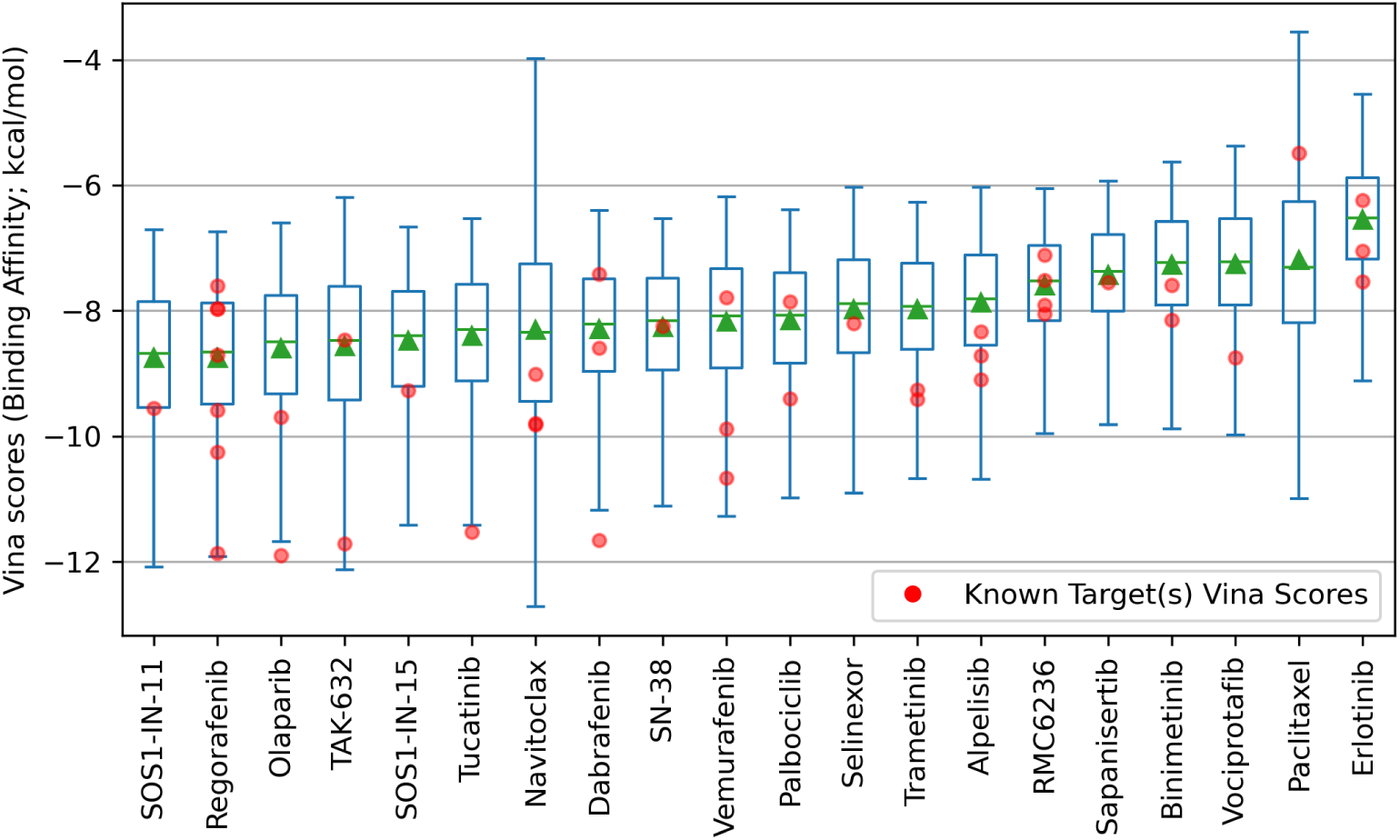
Plot visualizing the distribution of AutoDock Vina scores (Binding Affinity in kcal/mol) of docked complexes across 20 tested ligands and 7,657 gene-products selected for human proteome scan. Lower scores indicate stronger predicted binding affinities.

**Fig 5.**
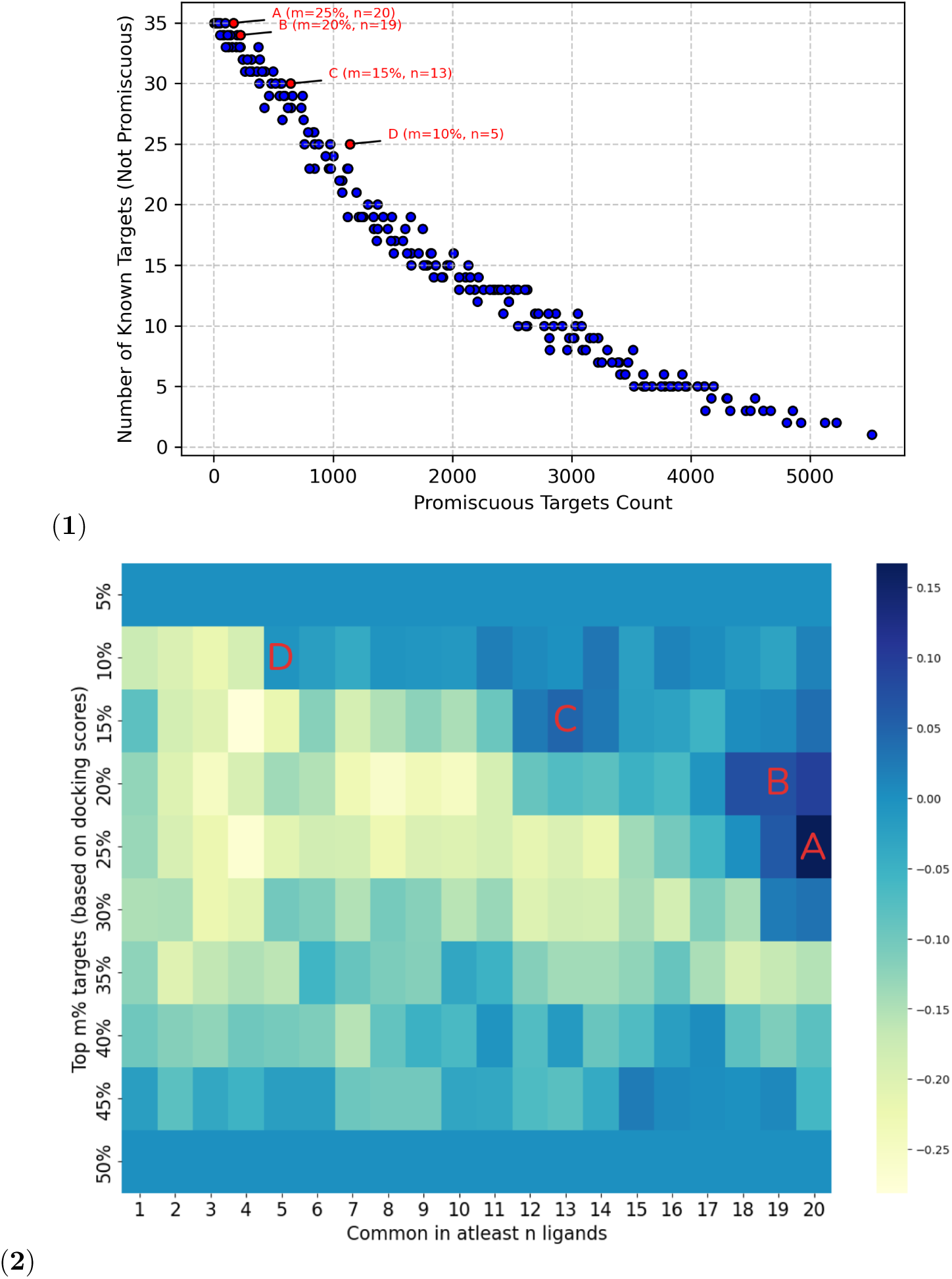
(1) Promiscuity Analysis (Across Thresholds): A) Thresholding the **top 25% targets common in all 20 ligands** gives 166 promiscuous targets and no known targets are found to be promiscuous. B) Threshold **top 20% targets common in at least 19 ligands** gives 222 promiscuous targets and 1 know target PARP1 is found promiscuous. C) Threshold **top 15% targets common in at least 13 ligands** gives 643 promiscuous targets and 5 know targets (HRAS, BRAF, BRAF V600E, PARP1, ERBB2) are found promiscuous. D) Threshold **top 10% targets common in at least 5 ligands** gives 1139 promiscuous targets and 10 know targets (HRAS, BRAF, PIK3CA E545K, KRAS, RET, BRAF V600E, PARP1, FLT1, PARP2, ERBB2) are found promiscuous. (2) Heatmap shows the difference between all promiscuous targets (min-max normalized scores) and known targets among promiscuous ones (min-max normalized) across varying thresholds. The x-axis represents the minimum number of ligands a target is common in, while the y-axis denotes the top m% of targets selected based on docking scores. The threshold cells, highlighted in darker blue, have higher number promiscuous targets with lesser number of known targets categorized as promiscuous.

**Fig 6.**
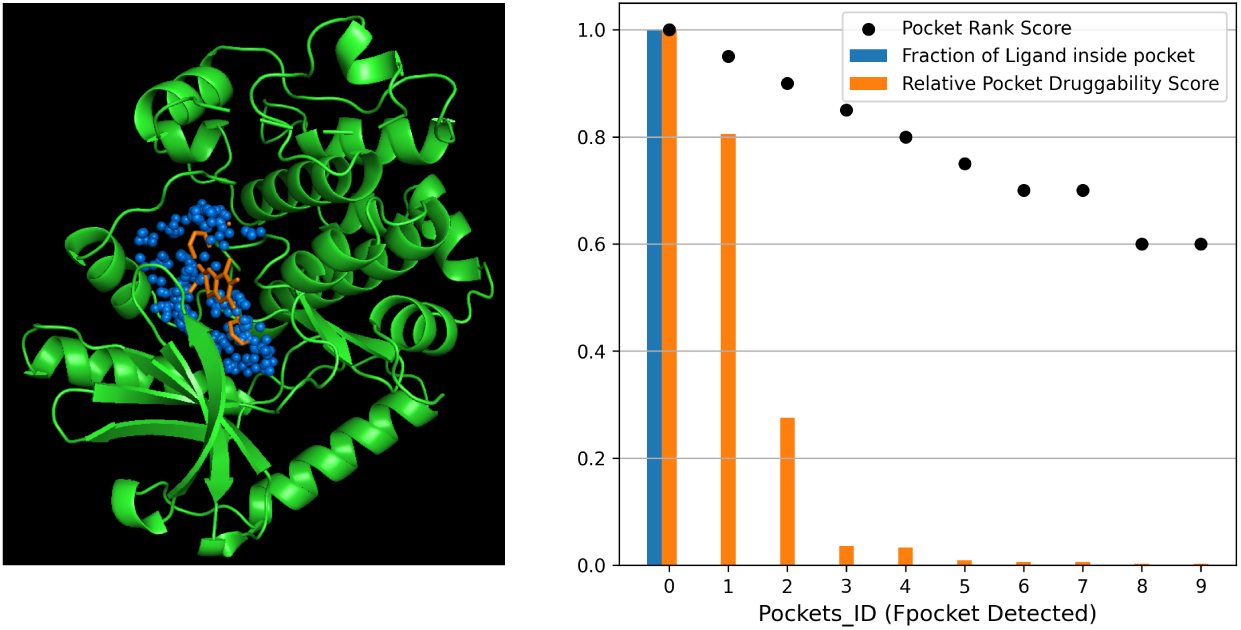
Binding pose analysis for MAP2K1-Trametinib complex. The image on the left represents binding pose of MAP2K1 (PDB ID 7M0X, chain B) with Trametinib. The plot on the right shows that the Trametinib ligand is 100% inside the pocket in MAP2K1-7M0X protein with highest druggability rank based on fpocket analysis, which validates a positive control of biologically relevant binding.

**Table 2.** Custom ligand-pocket interaction metrics used in pose evaluation.

| Metric Name | Description |
| --- | --- |
| % Pocket Filled | Percentage of alpha-spheres in a given pocket that are within 3 Å of any ligand atom. Serves as a proxy for how well the ligand fills a pocket. |
| % Ligand Inside Pocket | Percentage of ligand atoms that are within 3 Å of any alpha-sphere in the pocket. Measures how deeply the ligand is situated within a pocket. |
| Total % Ligand Inside Pocket | Sum of % Ligand Inside Pocket values across all detected pockets in the protein. Indicates overall pocket engagement. |
| Total % Ligand Inside Top-n Pockets | Similar to above, but restricted to the top-*n* pockets ranked by Druggability Score. Focuses the analysis on the most druggable regions. |

## 3 Results

In this section, we present the key findings from the ProteomeScan docking experiments and the subsequent analysis of known targets, promiscuous targets, and non-promiscuous off-targets using the methods described above.

### 3.1 Docking Score Distributions and Promiscuity Analysis

Following the data preparation steps, we executed docking on the cleaned optimal PDB models corresponding to 7,657 unique genes against the 20 selected ligands, resulting in an overall average completion rate of 93.65% for over 300,000 docking tasks. Further information on configuration settings and computational failure modes is available in appendix section 6.1. The plot in figure 4 shows the distribution of docking scores, determined by AutoDock Vina, highlighting differences in binding affinity ranges and variability for each ligand across the proteome.

ProteomeScan provides us with top-ranked targets for each ligand. To refine the potential hits obtained from the full docking-based ranking, we filtered out targets that appeared to be promiscuous and capable of binding multiple ligands. Figure 5 summarizes our promiscuity analysis, presenting thresholds from table 3. These thresholds were chosen to minimize the number of known targets that were not classified as promiscuous, while maximizing the identification of promiscuous targets. The table in figure 11 in the Appendix highlights how the removal of promiscuous targets affected the relative ranking of known targets for various ligands. The ranks of known targets improved substantially when moderately filtering promiscuous targets, though aggressive filtering occasionally eliminated relevant known targets.

**Table 3.** The table shows the resulting number of promiscuous targets and number of known targets (categorized as promiscuous) at different *m* (targets in top *m*%) and *n* (target common in atleast *n* ligands) thresholds.

| $m$ | $n$ | Promiscuous Targets Count | Number of Known Targets (Promiscuous) |
| --- | --- | --- | --- |
| 25 | 20 | 166 | 0 |
| 20 | 19 | 222 | 1 |
| 15 | 13 | 643 | 5 |
| 10 | 5 | 1139 | 10 |

### 3.2 Performance of Known Targets

Having established the overall docking performance and score distributions across the proteome, we next evaluated how well ProteomeScan recapitulates known ligand–target relationships (from table 1). This assessment provides a baseline for interpreting downstream predictions and validating the robustness of the docking pipeline.

#### 3.2.1 Known Target Recovery

To evaluate ProteomeScan’s known target prioritization performance, we define a coverage-level metric termed *Known Target Recovery* (KTR).

Let *N* denote the total number of known ligand-target pairs in the study (here ProteomeScan), and let *p_j_* denote the percentile rank of the known target associated with the *j*-th ligand-target pair within its corresponding proteome-wide ranking in ProteomeScan results. For a given cutoff *m*%, KTR quantifies the fraction of known interactions that are recovered within the top *m*% of the targets ranked.

Formally, KTR at cutoff *m*% is defined as

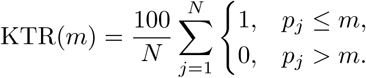

KTR was evaluated at multiple coverage thresholds (*m* = 5, 10, 15, and 30). This analysis was repeated with various levels of promiscuous targets (166, 222, 643, 1139) removed before ranking. As a baseline, we generated random rankings by assigning scores at random and selecting the top *m*% using the same thresholds. This random control was repeated across three different random seeds to account for stochastic variation. The ProteomeScan results outperformed the random baseline as shown in table 4.

**Table 4.** This table shows KTR(*m*) values (percentage of known targets (of the 20 drugs) retrieved within the top *m*% of ranked predictions), with *m* values of 5, 10, 15, and 30, for random baseline (averaged across 3 runs), ProteomeScan full run and ProteomeScan with various levels of promiscuous targets (166, 222, 643, 1139) removed before ranking. This table also shows Mann-Whitney U test p-values between random baseline and ProteomeScan full run and with various levels of promiscuous targets removed before ranking, respectively.

| Method | Top 5% | Top 10% | Top 15% | Top 30% | p-value |
| --- | --- | --- | --- | --- | --- |
| Random Baseline | 3.033 ± 2.627 | 9.850 ± 5.718 | 14.393 ± 8.600 | 34.090 ± 8.194 | - |
| ProteomeScan | 13.64 | 25.00 | 31.82 | 56.82 | 0.000063 |
| ProteomeScan (166 Promiscuous) | 20.45 | 29.55 | 36.36 | 59.09 | 0.000019 |
| ProteomeScan (222 Promiscuous) | 18.18 | 27.27 | 34.09 | 56.82 | 0.000025 |
| ProteomeScan (643 Promiscuous) | 15.91 | 22.73 | 34.09 | 47.73 | 0.000132 |
| ProteomeScan (1139 Promiscuous) | 13.64 | 25.00 | 27.27 | 40.91 | 0.000259 |

Mann-Whitney U tests on the ranks of known targets show that the observed differences between the ProteomeScan results and the random baseline are statistically significant. The p-values are also shown in table 4,

#### 3.2.2 Pose Analysis of Known Protein-Ligand Complexes

To further validate the ProteomeScan docking results, we applied the pose-analysis pipeline to known protein–ligand complexes (e.g., Figure 6). As a check on the pipeline’s accuracy, we found that the pocket-occupancy metrics aligned well with visually verified poses. Ligands occupying meaningful binding pockets exhibited high *% Ligand Inside Pocket* values, particularly within the top five pockets ranked by Druggability Score. These observations support the use of our metrics for identifying potentially (biologically) relevant binding modes.

We also noted that including all detected pockets can introduce noise, as many correspond to shallow or low-quality surface features, whereas restricting the analysis to only the top-ranked pocket by druggability can be overly narrow—especially for ligands spanning multiple adjacent pockets. Our results indicate that evaluating the top 5–10 pockets provides an effective balance, capturing relevant interactions while avoiding dilution from irrelevant pockets.

Based on the top 5–10 pocket analysis shown in Figure 7, we observe that the majority of known protein–ligand complexes exhibit *% Ligand Inside Pocket* values above 50%, consistent with expected binding poses reported in the literature. Limitations for complexes with lower *% Ligand Inside Pocket* scores are discussed in the following sections.

**Fig 7.**
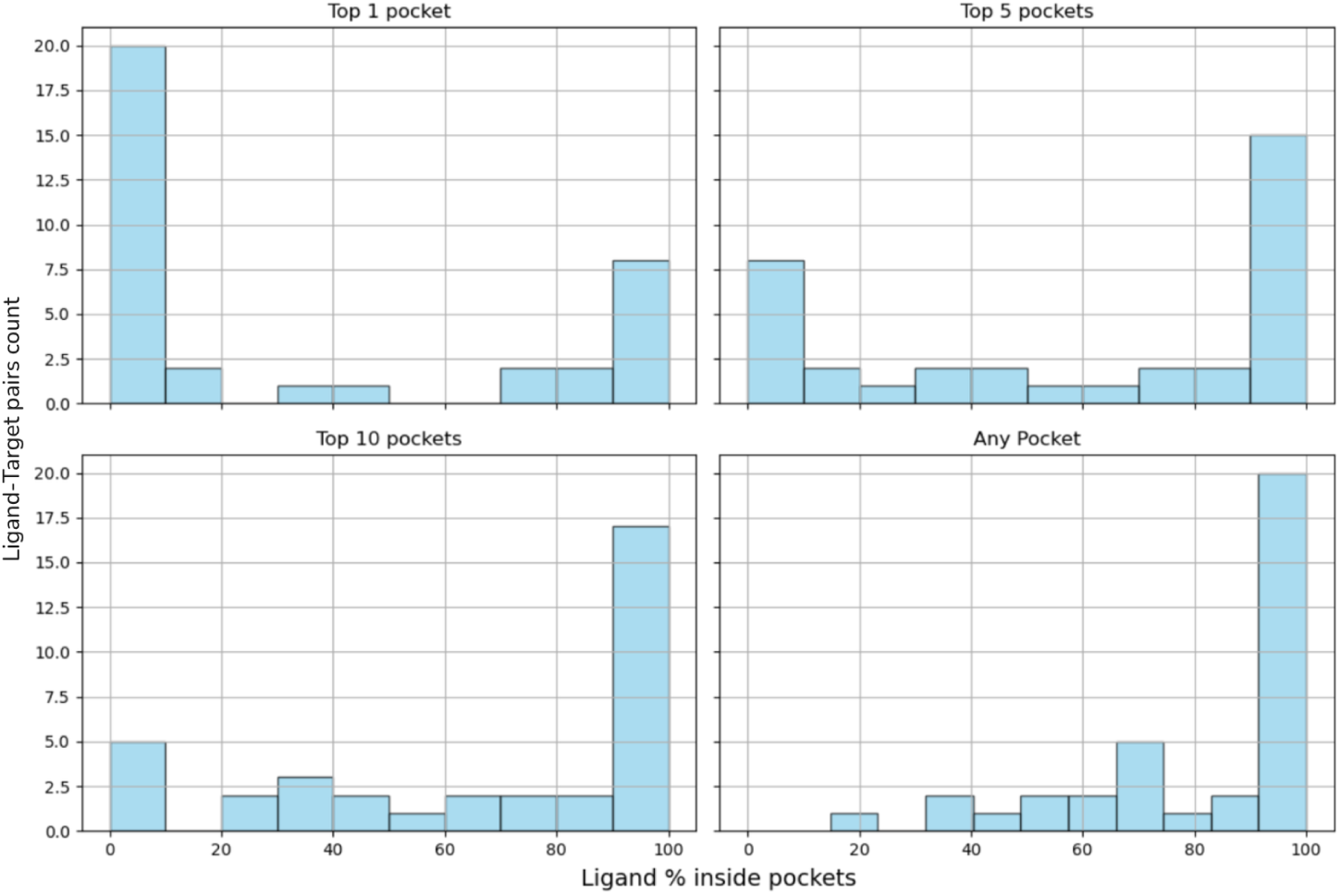
Pocket Occupancy analysis of ligands in their respective known targets.

#### 3.2.3 Mutants Analysis

We also evaluated ProteomeScan’s performance on well characterized drug mutation pairs where clinical evidence supports enhanced drug binding. This validation tests whether ProteomeScan can correctly predict that specific mutant variants should rank better than their wild-type counterparts as supported by experimental data. The specific results for mutants are summarized in the table 5.

**Table 5.**
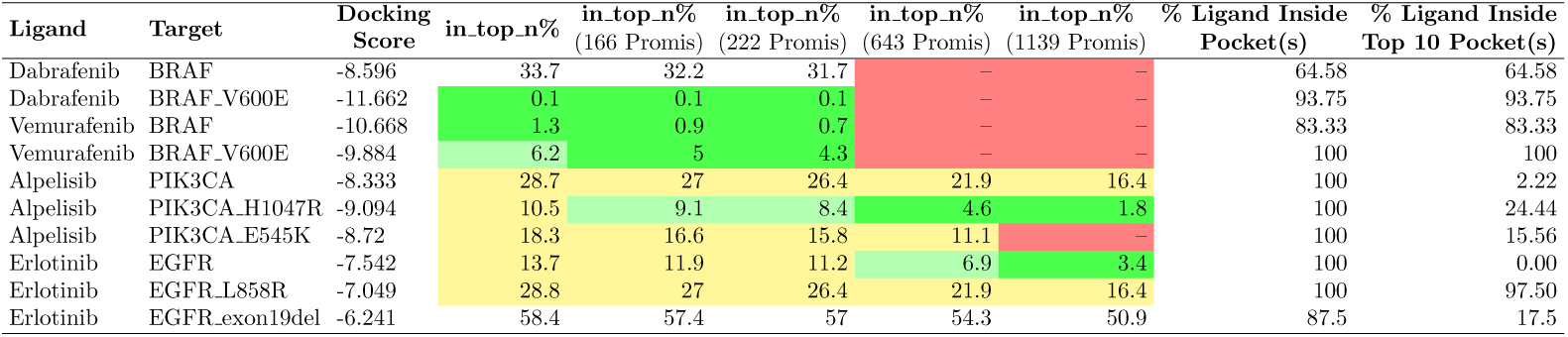
This table presents a comparative analysis of wild-type targets and mutant variants. Legends: 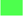: ≤ *top* 5%,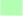: ≤ *top* 10%,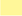: ≤ *top* 30% and 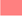: *target as promiscuous*.

Our analysis demonstrates the successful identification of clinically relevant drug-mutation interactions for specific protein families. For BRAF mutations,

ProteomeScan correctly predicted that BRAF V600E would rank better than wild-type BRAF with dabrafenib (rank 6 vs. 2459), aligning with clinical which has been detailed in the case study in 6.3.1. This prediction matches the mechanism where dabrafenib preferentially binds to the active conformation of BRAF kinase through competitive occupation of the ATP binding pocket, with confirmed activity against V600E mutations [32]. Similarly, ProteomeScan successfully predicted PIK3CA mutations, with H1047R ranking better than E545K, both superior to wild-type (ranks 759 and 1322 vs 2066 respectively). This matches preclinical data showing both H1047R and E545K mutations are potently inhibited by alpelisib, with H1047R in the kinase domain demonstrating stronger sensitivity than E545K in the helical domain [33].

ProteomeScan showed predictions for other established drug-mutation pairs, with vemurafenib-BRAF V600E ranking poorly despite clinical evidence of selective efficacy in V600E-positive melanomas [34], and EGFR mutations (L858R, exon19del) ranking worse than wild-type with erlotinib, contradicting their established role as primary FDA-approved biomarkers with higher binding affinity than wild-type EGFR [35]. The pocket analysis reveals critical explanations for these discrepancies. Successful predictions (BRAF V600E with dabrafenib, PIK3CA H1047R with alpelisib) showed high ligand occupancy in druggable pockets (93.75% and 100% respectively). In contrast, wild-type EGFR showed 0% ligand occupancy in the top 10 pockets, indicating erlotinib bound to an incorrect, non-druggable site, which explains its misleadingly favorable ranking. The EGFR mutants, despite their poorer rankings, actually showed much higher pocket occupancy (L858R: 97.50%, exon19del: 17.5%), suggesting they were binding in more relevant sites. Similarly, for vemurafenib-BRAF interactions, the pocket analysis reveals that wild-type BRAF (rank 90) showed 83.33% pocket occupancy, while the clinically relevant V600E mutant (rank 450) demonstrated perfect 100% pocket occupancy. This indicates that despite the poorer ranking, the V600E mutant actually achieved superior binding geometry in druggable pockets, aligning with clinical data. These mutant analyses demonstrate ProteomeScan’s capability to identify genuine high-affinity drug-target interactions while showing that binding scores alone can be misleading when ligands bind to irrelevant surface sites.

#### 3.2.4 Discussion on Complex Binding Requirements

In reviewing the ProteomeScan results for known targets, we observed cases with complex binding requirements that arise when drugs bind through mechanisms including allosteric conformational changes, and multimeric structure formation.

1. **Allosteric Mechanisms**: Several compounds in our test set bind through allosteric mechanisms (Table 6). Allosteric inhibitors bind to sites distinct from the active site and induce conformational changes in the target protein. With blind-docking based approach of ProteomeScan, allosteric inhibitors demonstrated favorable performance compared to non-allosteric compounds. The mean in top n% is 17.6% for allosteric compounds vs 28.9% for non-allosteric compounds (Figure 11, excluding multimeric structures that we discuss below). Because of the small number of allosteric and non-allosteric targets, this comparison should not be considered statistically rigorous. However, ProteomeScan’s results suggest that it may be able to model allosteric effects effectively. We have a detailed case study on Trametinib in Appendix 6.3.1, where we discuss allostery further.
2. **Multimeric Structure-Dependent Mechanism**: Our paclitaxel validation study (detailed in Appendix 6.3.3) illustrates computational challenges when drugs require binding to macromolecular assemblies rather than isolated protein structures. The known target TUBB ranked poorly at position 5724 of 7,657 targets, with docking scores of -5.485 (8V2J monomer) and -7.471 (8V2J multi-chain). This modest ranking reflects fundamental limitations in modeling assembly-dependent binding rather than a failure of the methodology itself.

**Table 6.** Ligand - known target pairs with allosteric site binding. Percentages indicate the ranking position (top *n*%) among all 7,657 targets

| Ligand | Target | In_top_n% |
| --- | --- | --- |
| Binimetinib | MAP2K2 | 15.7 |
|  | MAP2K1 | 33.1 |
| Trametinib | MAP2K2 | 6.2 |
|  | MAP2K1 | 8.5 |
| TAK-632 | BRAF | 0.9 |
|  | RAF1 | 46.3 |
| SOS1-IN-11 | SOS1 | 22.1 |
| SOS1-IN-15 | SOS1 | 20.4 |
| Vociprotafib | PTPN11 | 5.6 |

Paclitaxel’s binding mechanism requires assembled microtubules rather than free tubulin dimers. The taxane binding site is occluded by *β*M-loop dynamics in unassembled tubulin, rendering it inaccessible in isolated dimers [36]. This assembly dependence manifests as a 100–1000 fold affinity difference: paclitaxel binds assembled microtubules with *K_D_* ∼10–50 nM [37] versus free tubulin dimers at *K_D_* ∼1–10 *µ*M [38].

Similarly, RMC-6236 exhibited poor computational ranking for PPIA (top 61.2% among 7,657 targets) as it requires formation of an inhibitory tri-complex between cyclophilin A (PPIA), the inhibitor, and RAS proteins [39]. The inhibitor acts as molecular glue to induce a non-native PPIA:RAS interaction that blocks RAS signaling, a mechanism that cannot be captured by docking to individual protein structures. Static docking algorithms operating under rigid receptor assumptions cannot model the large-scale conformational reorganization and cooperative binding energetics required for high-affinity paclitaxel-tubulin interactions. This limitation is well-documented for assembly-dependent binding systems [40], where experimental validation remains essential.

### 3.3 Further Characterization of Promiscuous Targets

As summarized in Table 3, varying the threshold yields different sets of promiscuous targets. Using a threshold defined by the top 25% of targets shared across all 20 ligands results in a set of 166 promiscuous targets, none of which overlap with the known targets. In the sections that follow, we examine these 166 targets in greater detail (the complete list is provided in Appendix 6.5).

#### 3.3.1 Biological Basis of Promiscuous Target Identification

Our systematic identification of 166 promiscuous targets provides valuable insights into the molecular basis of broad small-molecule binding across the human proteome. The majority of identified targets belong to well-characterized protein families with documented ligand promiscuity, validating ProteomeScan’s ability to identify biologically relevant binding patterns, while others represent candidates requiring experimental validation.

Metabolic enzymes with documented promiscuity are the most strongly supported category, including drug-metabolizing enzymes. CYP3A4 emerged as promiscuous, consistent with its role in metabolizing approximately 50% of marketed drugs due to its large, flexible active site [41, 42]. Butyrylcholinesterase (BCHE) appeared in our list, aligning with its established function as a biological scavenger that hydrolyzes diverse esters and neutralizes toxic compounds [43]. Glutathione S-transferase GSTA1 demonstrated promiscuous binding, matching previous findings that GST isoforms exhibit broad catalytic promiscuity due to their role in xenobiotic detoxification [44].

Heat Shock Proteins such as HSP90AA1 were identified as promiscuous, consistent with literature documenting multiple inhibitor chemotypes binding to its ATP pocket [45, 46]. Aldo-Keto Reductases similarly exhibit broad substrate specificity enabling Phase I metabolism of diverse pharmaceuticals and xenobiotics [47]. Multiple AKR family members (AKR1B1, AKR1B10, AKR1C3) appeared as promiscuous targets, supported by structural studies demonstrating broad substrate specificity due to their role in phase I metabolism [48].

Alcohol and Aldehyde Dehydrogenases were another well-documented category. Both ADH family members (ADH4, ADH5) and ALDH family enzymes were identified as promiscuous. Literature supports the broad substrate scope of these enzyme families, with ADHs contributing to diverse xenobiotic metabolism beyond ethanol oxidation [49], and ALDHs handling both endogenous and exogenous aldehyde substrates [50].

Additional promiscuous targets include HTR2B (5-HT2B receptor), a well-characterized anti-target where agonist activity has led to drug withdrawals due to cardiac valvulopathy, prompting routine screening in drug discovery [51]. SLC transporters (SLC14A1, SLC2A3, SLC9A3) showed promiscuity characteristic of proteins mediating drug absorption and disposition [52]. VKORC1 demonstrated promiscuity consistent with being targeted by structurally diverse anticoagulants [53], while ACAD9 and AOX1 belong to enzyme families with broad substrate specificity in fatty acid metabolism and drug biotransformation [54, 55]. KDM1A exhibited binding to diverse inhibitor scaffolds reflecting its therapeutic relevance across multiple indications [56].

Several identified targets (HDAC8, SCN3A, CHRM1, DYRK2) possess large or highly flexible binding sites that may lead to over-prediction in docking-based approaches [57]. Experimental validation of these targets is necessary to distinguish genuine polypharmacology from computational artifacts arising from permissive binding pockets.

#### 3.3.2 Pose Analysis of Promiscuous Targets

For the 166 promiscuous targets, it remains unclear whether the ligands are engaging biologically relevant binding pockets, or if they are simply located near surface-exposed regions, producing artificially high docking scores without meaningful binding. As shown in figure 8(A), pocket occupancy analysis of around 3300 complexes with these promiscuous targets reveals that a majority of the ligand–target complexes exhibit high *% Ligand Inside Pocket* values, particularly within the top five pockets ranked by fpocket druggability scores. However, a small subset of complexes also display ligand binding outside of these predicted pockets, indicating potential out-of-pocket interactions.

**Fig 8.**
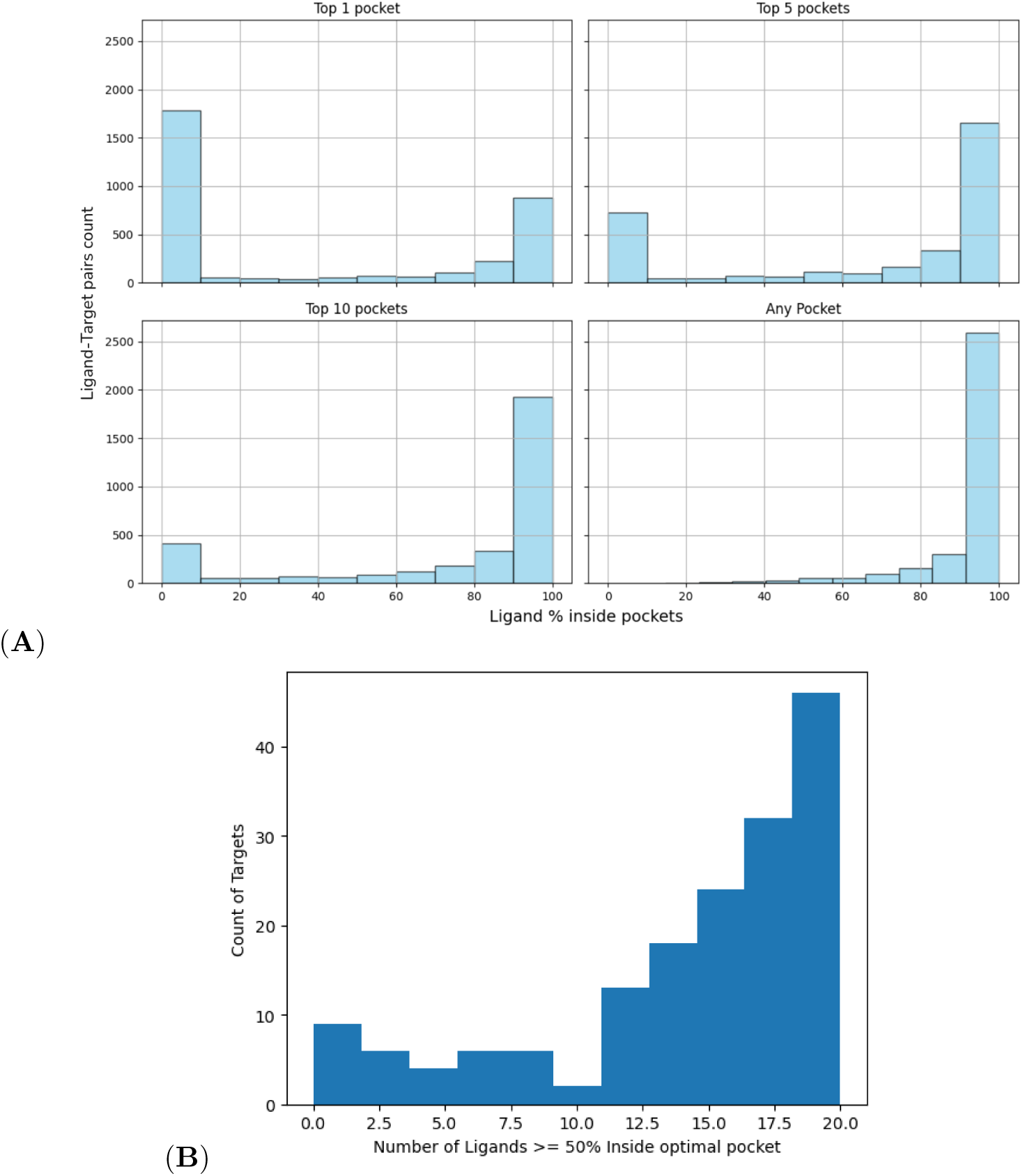
A total of 166 promiscuous targets appeared in the top 25% of docking scores across all 20 ligands. (A) Pocket Occupancy analysis of Ligands in 166 Promiscuous Targets. The unit of count in y-axis is based on total of 3320 unique ligand-target pairs from 20 ligands and 166 promiscuous targets. (B) For each of these promiscuous targets, we evaluated how many ligands are located within the top-5 pockets predicted by fpocket. A ligand is considered to bind a druggable pocket if more than 50% of its atoms fall within these pockets. The results suggest that most promiscuous interactions occur within druggable pockets.

For each of these promiscuous targets, we evaluated how many ligands were located within the top five pockets predicted by fpocket, shown in figure 8(B). A ligand was considered to bind a druggable pocket if more than 50% of its atoms were contained within these regions. The results indicate that the majority of promiscuous interactions involve druggable pockets, supporting the retention of these targets as promiscuous for downstream analysis and filtering.

#### 3.3.3 Methylation Analysis

As an alternative approach to assess whether high docking scores for promiscuous targets reflect a favorable binding pose, we investigated the impact of ligand methylation on binding affinity, focusing specifically on kinase-based promiscuous targets. Previous studies [58] have shown that the addition of methyl groups to drug molecules often reduces their affinity for kinase-binding pockets. Consequently, if methylation does not lead to a significant change in binding affinity, then it may suggest that the high negative docking scores observed are not indicative of biologically relevant kinase interactions.

Our docking analysis on a subset of kinase targets from the 166 promiscuous targets, using methylated ligands, is presented in Figure 9. In general, the results show a decrease in binding affinity for the methylated variants compared to their non-methylated counterparts. However, a few cases exhibit minimal change in docking scores following methylation, suggesting weak or non-specific interactions with the kinase targets in those instances.

**Fig 9.**
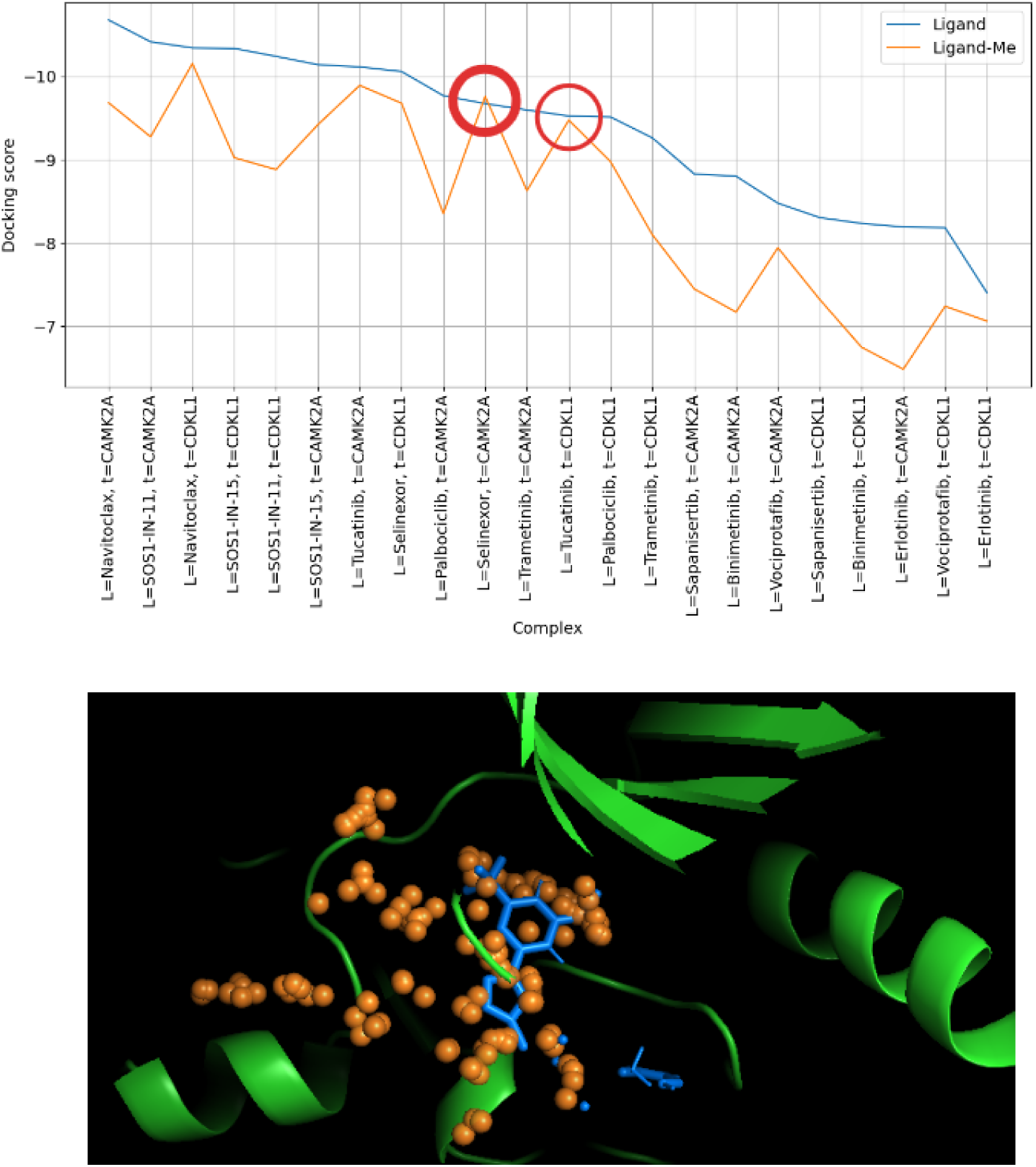
The plot on the top shows docking scores for a subset of kinase promiscuous targets complexes with original ligands (line in blue) and with methylated form of ligands (line in orange) in a sorted order. The bold highlighted red circle shows CAMK2A-Selinexor complex where docking score did not change significantly before and after methylation. The image on the bottom shows docked complex for CAMK2A (PDB ID 2VZ6, chain A/B) and methylated Selinexor. The orange spheres depict the druggable pocket identified by fpocket.

### 3.4 Non-promiscuous off-targets

While evaluating the ProteomeScan results, we identified potential off-targets through binding affinity patterns, promiscuity filtering, and pose analysis. For the three ligands studied in detail (see Appendix 6.3), the identified off-targets tended to be either homologous or functionally related to their known targets despite the functional connections being relatively weak. Notable observations include the case of Dabrafenib where we find a hydroxy-dabrafenib metabolite (with a hydroxyl group attached to dabrafenib) is known to inhibit CYP1A2 (IC_50_ = 83 *µ*M) [59]. To further validate this observation, we carried out an independent docking computation, which demonstrated favorable binding of hydroxy-dabrafenib to CYP1A2 (PDB ID: 2HI4), yielding a docking score of −10.91. This known related binding interaction supports our predicted interaction for Dabrafenib and CYP1A2, and suggests potential metabolic modulation by dabrafenib or its derivatives.

## 4 Discussion

Our results suggest that ProteomeScan can capture several meaningful trends in proteome-wide docking, though many findings should be interpreted with care. The docking score distributions and subsequent filtering allowed us to narrow an initial set of candidate targets by excluding those that appeared broadly promiscuous across ligands. Analyses of known targets showed statistically significant separation from random baselines and generally reasonable agreement with expected binding poses, although some cases exhibited lower pocket-occupancy values and highlight the limits of docking-only assessments. Additional evaluations on mutants and assembly-dependent or allosteric cases indicate that certain clinically relevant interactions can be recovered, but they also point to computational limitations of the docking methology used in ProteomeScan.

Our characterization of the 166 promiscuous targets suggests that many fall into protein families where broad ligand binding is biologically plausible, and pocket-level analyses provide some support for treating these as genuine promiscuous interactions. Follow-up analysis using methylated ligands further illustrate that differences in predicted affinity are not uniform across targets, reinforcing the need for cautious interpretation. Finally, after removing promiscuous proteins, a small number of non-promiscuous off-targets emerged for some ligands, aligning to varying degrees with homology or functional hints reported in the literature. Together, these observations indicate that ProteomeScan may help prioritize candidate targets, but currently high false-positive and false-negative rates suggest that ProteomeScan is best used alongside with additional target validation approaches.

### 4.1 Known Caveats

Several important caveats should be considered when interpreting the results of this study. Firstly, protein-protein interactions were not explicitly considered, which may influence docking accuracy, as interactions in vivo can alter available binding sites. Secondly, our docking analyses did not fully account for multimer-based dynamic pockets, potentially overlooking biologically relevant conformational changes.

Additionally, post-translational modifications such as phosphorylation, glycosylation, and acetylation were not incorporated into our protein structures, which could significantly affect binding site accessibility and drug-protein interactions in physiological conditions. Finally, as with any computational method, it remains challenging to precisely estimate the degree of error inherent in docking predictions, underscoring the necessity for experimental validation of computationally predicted interactions.

In terms of scalability, the overall percentage completion of the ProteomeScan could be further improved with better handling of spot instance interruptions. See section 6.1 in Appendix for more details.

### 4.2 Known Target Recovery as an Evaluation Metric for Docking Algorithms

ProteomeScan introduces a new evaluation metric, referred to as *Known Target Recovery (KTR)*, aimed at assessing the performance of docking functions at the proteome-level. In principle, this metric could be valuable for systematic evaluation and benchmarking of different deep docking approaches, as well as for testing and comparing newly developed methods in future studies. However, it is important to note that, in current formulation of ProteomeScan, the metric has limitations when applied to systems with complex binding modes such as heterogeneous interactions in allosteric complexes. Consequently, there is a risk that optimizing too aggressively for this metric could lead to models that poorly capture realistic binding behavior.

### 4.3 Applications in Drug Repurposing

Target identification has been foundational for drug repurposing [3], with early studies demonstrating how computational screens could identify unexpected targets for approved drugs. Drug repurposing studies have demonstrated methods to adapt existing drugs to new targets. For example, nitroxoline, originally an antibiotic, was identified through virtual screening as a selective BRD4 inhibitor for treating MLL leukemia [60], while berberine was found to target UHRF1 in multiple myeloma [61]. Similarly, pridopidine was expanded from Huntington’s disease to other neurodegenerative diseases after identification as a *σ*_1_ receptor agonist [62], and azidothymidine (AZT) was repurposed from chemotherapy to HIV treatment after structural studies revealed its mechanism against HIV reverse transcriptase [63]. These examples underscore the significance of target identification in drug repurposing by revealing previously unrecognized molecular interactions. ProteomeScan’s proteome-wide screening approach could accelerate drug repurposing by systematically identifying unexpected drug–target interactions across the human proteome. However, due to ProteomeScan’s high false-positive and false-negative rate, careful integration with complementary data sources would be needed to validate such results.

### 4.4 Potential Applications in Toxicology

Off-target toxicity causes over one-third of pharmaceutical development failures [64], often because companies typically test new drugs against limited panels of known problematic targets like hERG, CYP enzymes, or safety-related receptors [65]. This leaves thousands of proteins unexamined, creating blind spots where unexpected toxicities can emerge during expensive clinical trials. ProteomeScan could help address this by testing drug candidates against 7,500+ human proteins to identify potential off-target binding, particularly when integrated with complementary studies.

Kong et al. recently used proteome-wide docking to discover that PFAS chemicals bind to folate receptors, explaining their neurodevelopmental toxicity, a mechanism that would never have been found using traditional toxicology approaches focused on known targets [25]. This demonstrates how comprehensive protein screening can reveal unexpected toxicity pathways. By providing complete molecular interaction maps rather than limited target panels, future iterations of ProteomeScan could support more robust predictive safety assessments, especially once ProteomeScan’s currently high false-positive and false-negative rates are improved.

### 4.5 Deep Learning Scoring Functions

Recent advances in deep learning methods have shown promising improvements in molecular docking accuracy, but these methods face significant computational and methodological barriers for proteome-wide applications. Deep learning docking methods like GNINA 1.3 [66] and AlphaFold3 [67] face computational barriers for proteome-wide applications. Even GNINA 1.3’s fastest model is 1.3 seconds slower than Vina per docking calculation. More significantly, these methods exhibit training data biases [68] and provide confidence scores rather than interpretable binding energies, complicating ProteomeScan’s promiscuity analysis that requires consistent scoring behavior across diverse protein-ligand combinations. Additionally, deep learning approaches often fail to correctly identify native binding conformations despite high affinity prediction accuracy [69], and their performance varies significantly across different protein families [70]. Furthermore, machine learning-based scoring functions face the “black box” problem, where predictions cannot be easily interpreted or validated [71].

We selected AutoDock Vina for its computational scalability, methodological consistency across protein families [72, 73], and interpretable empirical scoring [74] essential for ProteomeScan’s systematic target identification and pose validation workflows. This approach enabled the large-scale blind docking required for systematic proteome scanning while maintaining reliable statistical comparisons across our diverse dataset. The scoring function’s transparent energy components facilitate the promiscuity analysis central to ProteomeScan’s methodology [75]. Future iterations could leverage deep learning methods for targeted refinement of high-priority predictions identified through Vina-based screening [76].

### 4.6 Open Source Availability

We have open sourced the code for ProteomeScan on https://github.com/deepforestsci/ProteomeScan, to promote transparency, reproducibility, and collaborative development within the research community.

## 5 Conclusion

A major challenge in drug discovery is that many early screens are phenotypic, testing compounds for a desired effect in cells or organisms without knowing which proteins they hit. This makes it difficult to pinpoint true targets, predict off-target liabilities, or rationally optimize efficacy and safety. By systematically docking across the entire human proteome, ProteomeScan moves beyond phenotype alone, revealing both on and off-targets before costly in vitro or in vivo experiments. This capability accelerates hit-to-lead progression, guides medicinal chemistry toward the most promising binding sites, and flags potential toxicity issues early.

## 6 Appendix

### 6.1 Docking Compute Details and Limitations

We configured AutoDock Vina with two key parameters:

- **Exhaustiveness = 32**: This setting determines the amount of computational effort spent in exploring the ligand–protein binding space. A higher exhaustiveness value significantly improves the robustness of the search by reducing the chance of missing low-energy binding poses in blind docking.
- **Number of modes = 8**: This parameter controls the number of top-ranked binding poses returned for each docking run. By retaining multiple plausible binding conformations, we increased the likelihood of identifying alternative binding modes that could be biologically relevant.

For the full ProteomeScan across 20 ligands and 7657 Genes (with 2 optimally selected PDB files per Gene on average as shown in figure 10), 300K+ docking runs were needed. These runs were parallely executed on Prithvi via AWS Spot instances with an overall average completion rate of 93.65%. Per-ligand completion rate can be checked in table 7.

**Fig 10.**
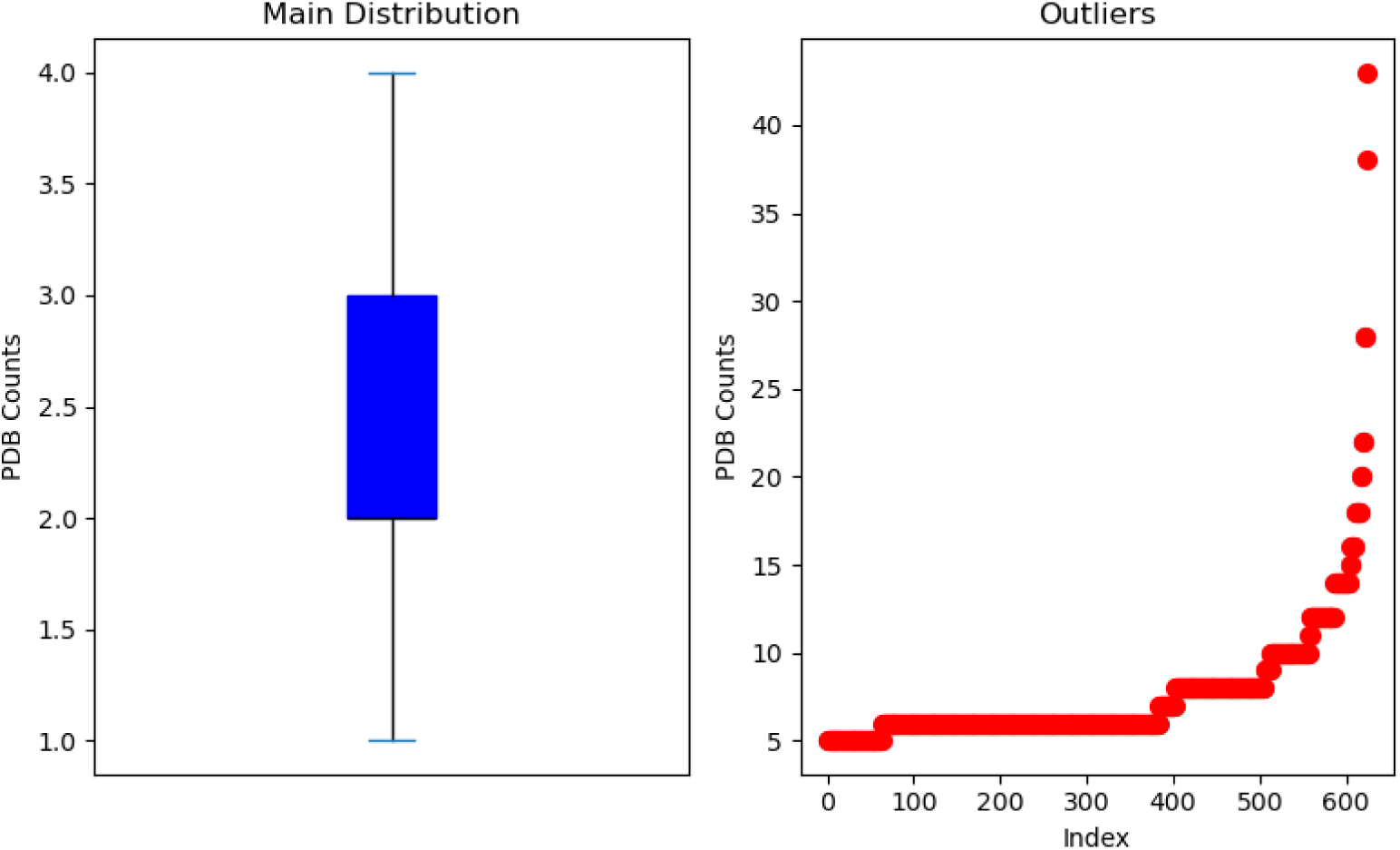
Distribution of number of optimally selected PDB files per Gene

**Table 7.** This table shows completion rate (out of 7657 Genes) of ProteomeScan across different ligands.

| Ligands | Gene Runs Completed | Completion Rate |
| --- | --- | --- |
| Sapanisertib | 7419 | 96.9 |
| SOS1-IN-15 | 7363 | 96.2 |
| Trametinib | 7333 | 95.8 |
| SOS1-IN-11 | 7330 | 95.7 |
| SN-38 | 7303 | 95.4 |
| Binimetinib | 7297 | 95.3 |
| Dabrafenib | 7288 | 95.2 |
| Regorafenib | 7278 | 95.1 |
| Palbociclib | 7273 | 95 |
| Tucatinib | 7270 | 94.9 |
| RMC6236 | 7266 | 94.9 |
| Vemurafenib | 7211 | 94.2 |
| Alpelisib | 7207 | 94.1 |
| Olaparib | 7202 | 94.1 |
| Selinexor | 7150 | 93.4 |
| TAK-632 | 7133 | 93.2 |
| Vociprotafib | 7094 | 92.6 |
| Erlotinib | 6866 | 89.7 |
| Paclitaxel | 6700 | 87.5 |
| Navitoclax | 6547 | 85.5 |

**Fig 11.**
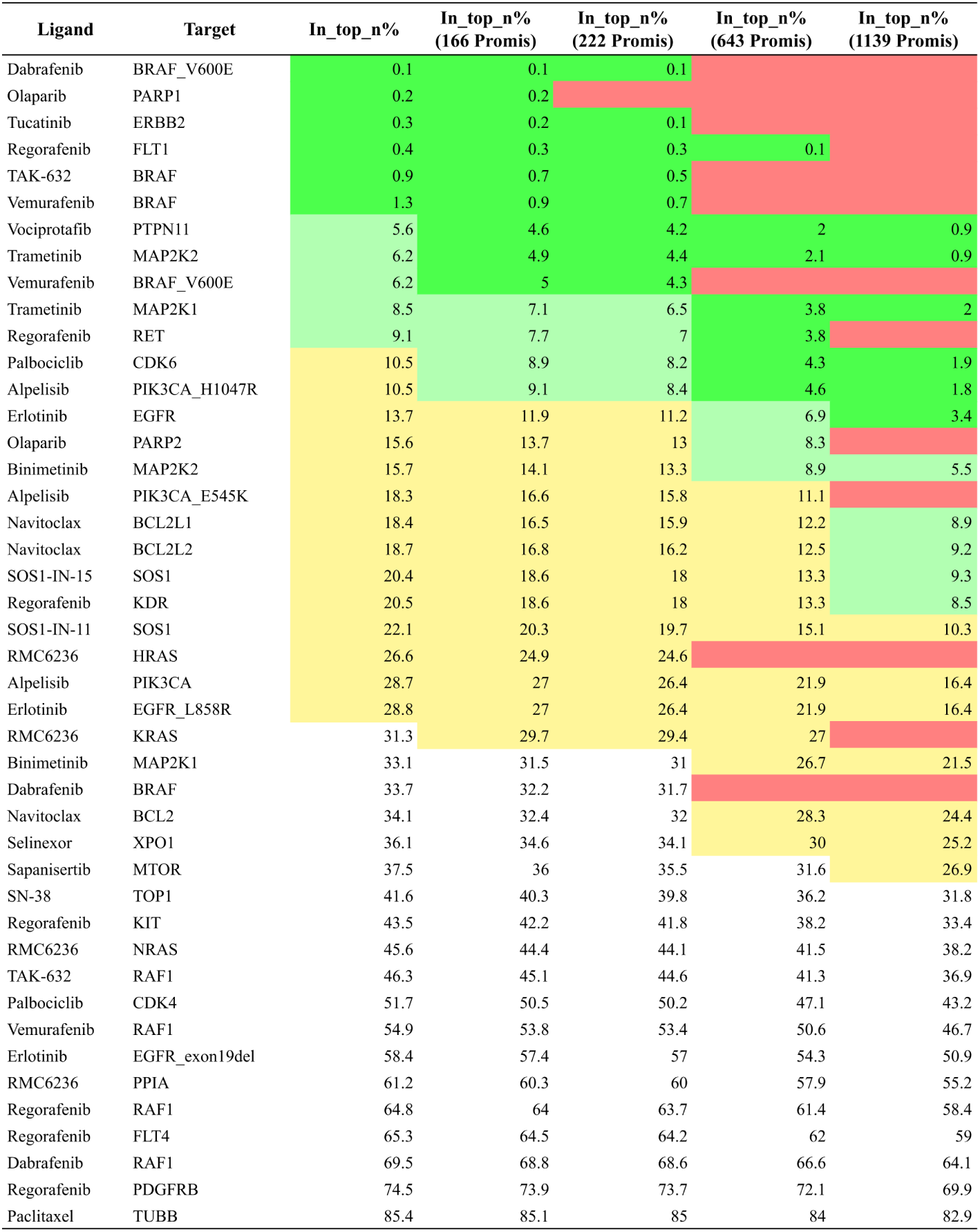
Results of promiscuity analysis. Legends: 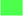: ≤ *top* 5%,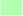: ≤ *top* 10%,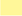: ≤ *top* 30% and 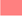: *target as promiscuous*.

**Fig 12.**
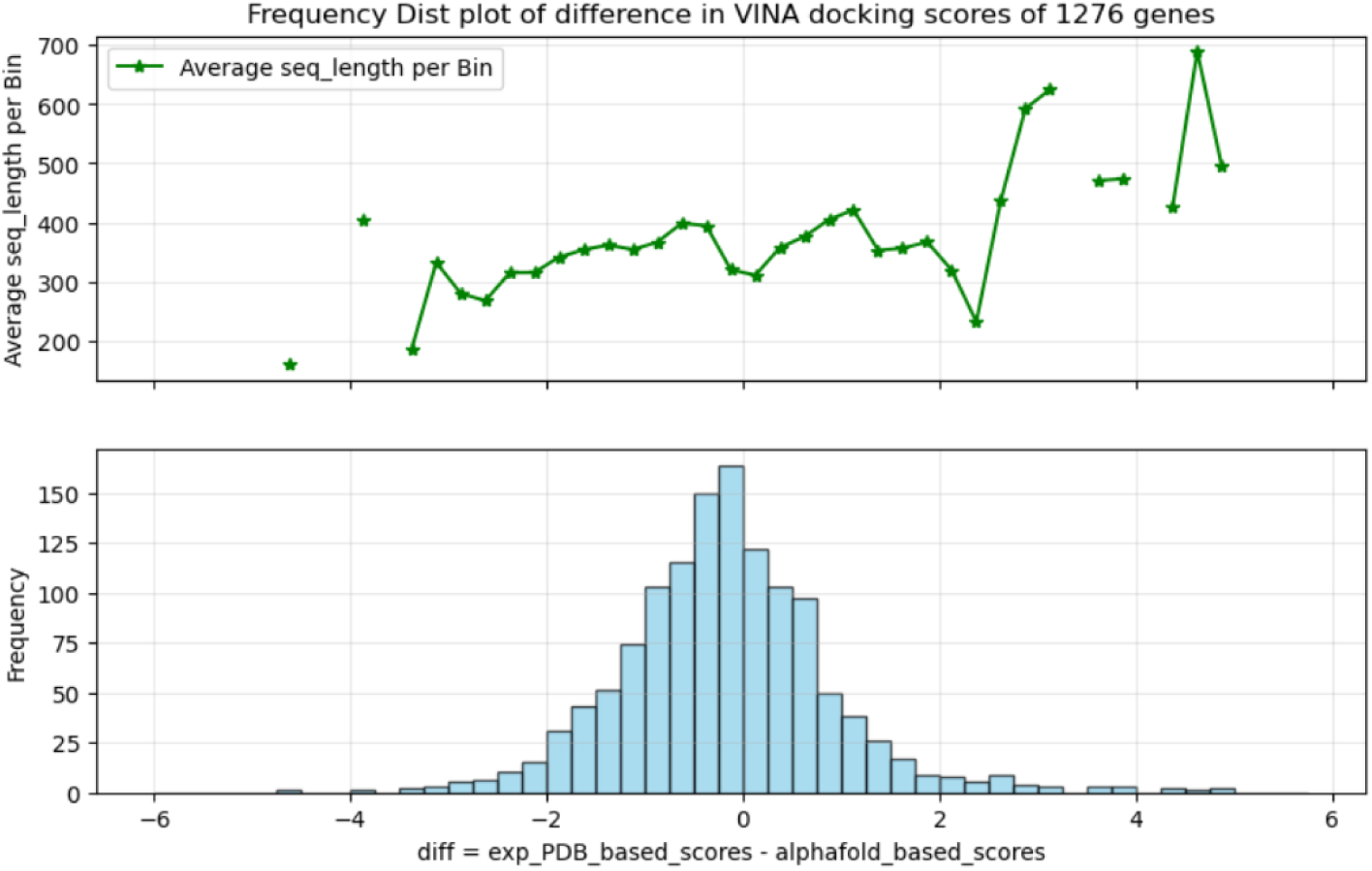
Trametinib Alphafold comparitive analysis

**Fig 13.**
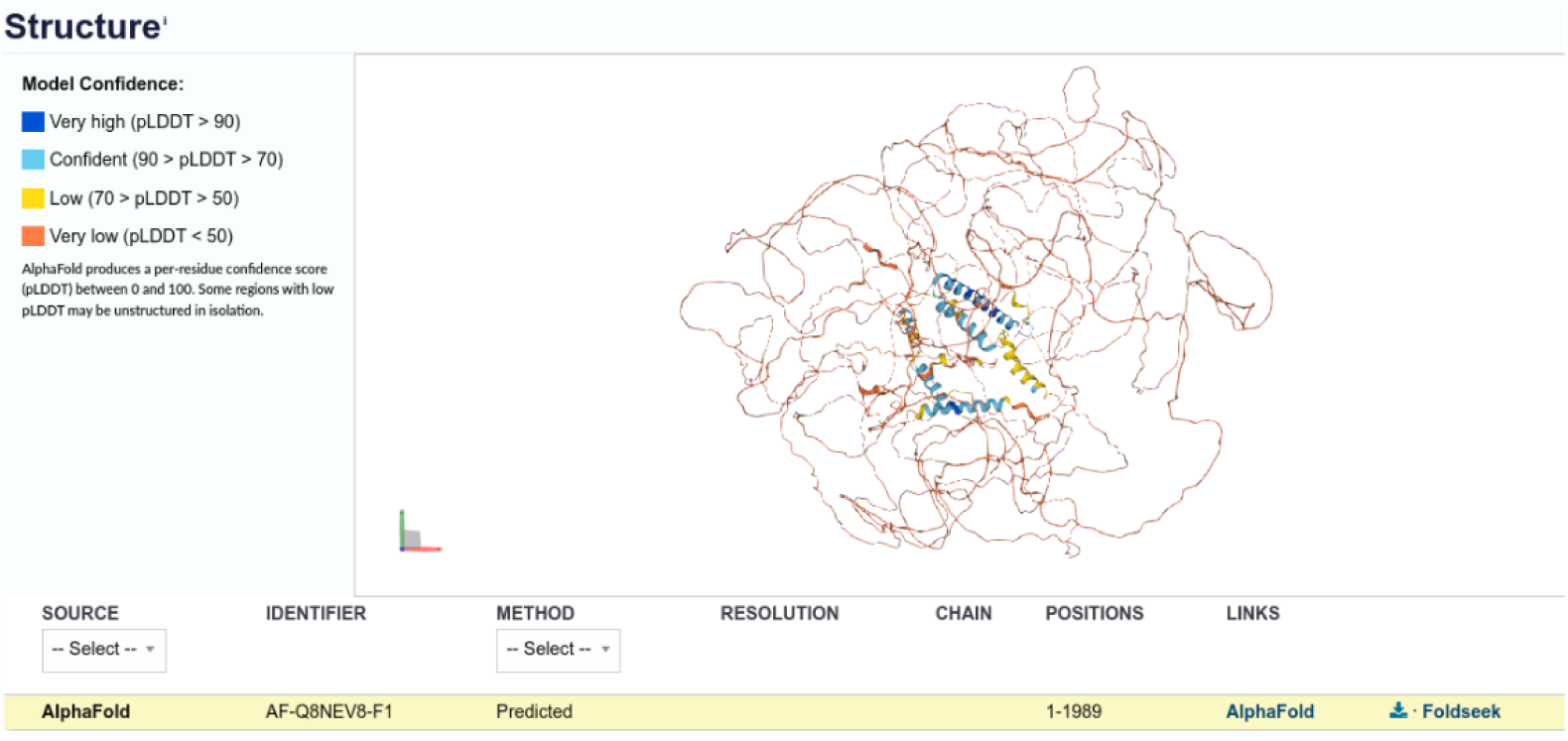
EXPH5 (PDB id: Q8NEV8) Alphafold Structure

One of the key reasons for failures in the docking runs was spot node interruptions, which were particularly evident in around 600 outlier cases involving genes with a higher number of PDB files to dock per gene (as shown in figure 10), since each spot node was responsible for handling all the docking runs for a given gene.

### 6.2 Promiscuity Analysis: Extended

#### 6.3 Case Studies

To demonstrate ProteomeScan’s capabilities and limitations in real-world scenarios, we present detailed analyses of three drug-target interactions. Starting with 7,657 unique genes, we performed proteome-wide docking and then applied promiscuity thresholds. Subsequently, we conducted pose analysis using fpocket on the top 10 targets, retaining only complexes where more than 50% of ligand atoms resided within druggable pockets as ranked by fpocket scores. Through the examples of dabrafenib, trametinib and paclitaxel, we demonstrate successful identification of known targets, highlight methodological limitations associated with conformationally complex binding mechanisms, and compare computational results with experimental literature to evaluate the biological relevance of predicted interactions.

##### 6.3.1 Case Study 1: Dabrafenib

This case study demonstrates ProteomeScan’s successful identification of clinically relevant off-targets for the BRAF inhibitor dabrafenib. In the initial target-identification scan, the known target BRAF V600E ranked 6th out of 7,657 targets. Its rank improved to 3 after applying the 25%(20) promiscuity filter, which excluded 166 targets, and pose filter.

###### Target Analysis

The identified targets (see Table 8) were categorized into three groups: (1) the validated on-target (BRAF V600E), (2) literature supported off-targets, and (3) computational predictions without experimental validation.

**Table 8.**
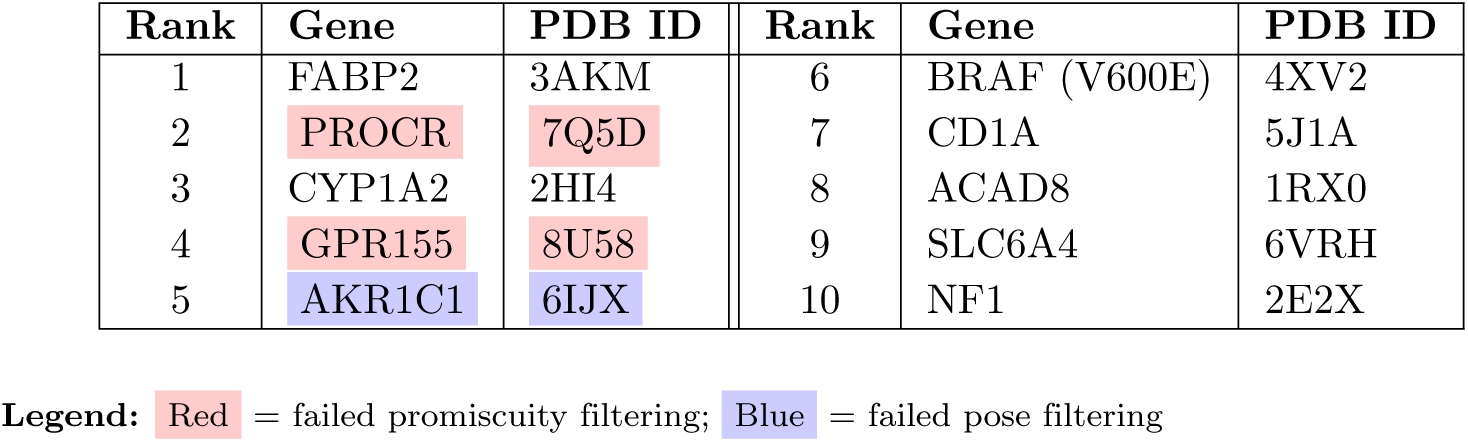
Dabrafenib off-targets identified after applying 25%(20) promiscuity filtering and pocket analysis validation. Promiscuity filtering removes proteins that appear in the top 25% of docking scores for all 20 ligands tested, representing the most broadly binding targets. Red highlighting indicates targets eliminated by promiscuity filtering, while blue highlighting indicates targets failing pose filtering.

| Rank | Gene | PDB ID | Rank | Gene | PDB ID |
| --- | --- | --- | --- | --- | --- |
| 1 | FABP2 | 3AKM | 6 | BRAF (V600E) | 4XV2 |
| 2 | PROCR | 7Q5D | 7 | CD1A | 5J1A |
| 3 | CYP1A2 | 2HI4 | 8 | ACAD8 | 1RX0 |
| 4 | GPR155 | 8U58 | 9 | SLC6A4 | 6VRH |
| 5 | AKR1C1 | 6IJX | 10 | NF1 | 2E2X |

**Legend:** Red = failed promiscuity filtering; Blue = failed pose filtering

**Table 9.**
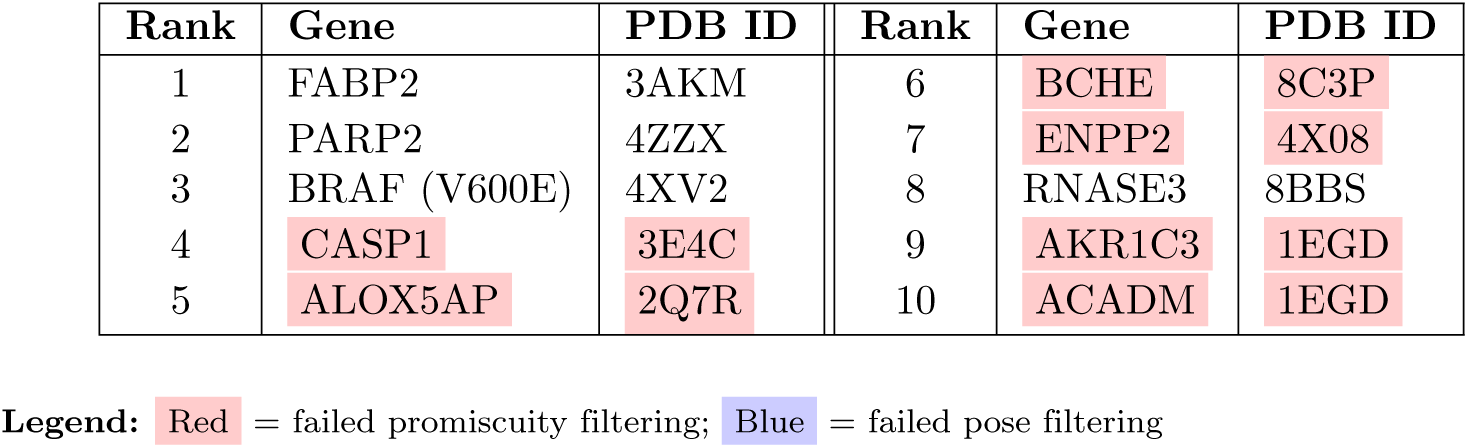
This table presents potential Trametinib off-targets,identified after a 25% promiscuity filtering (all 20 ligands) and pose filtering. This threshold removes proteins appearing in the top 25% of docking scores across all 20 ligands tested. Red highlighting indicates targets removed by promiscuity filtering and blue highlighting indicates failed pose filtering validation.

**Table 10.**
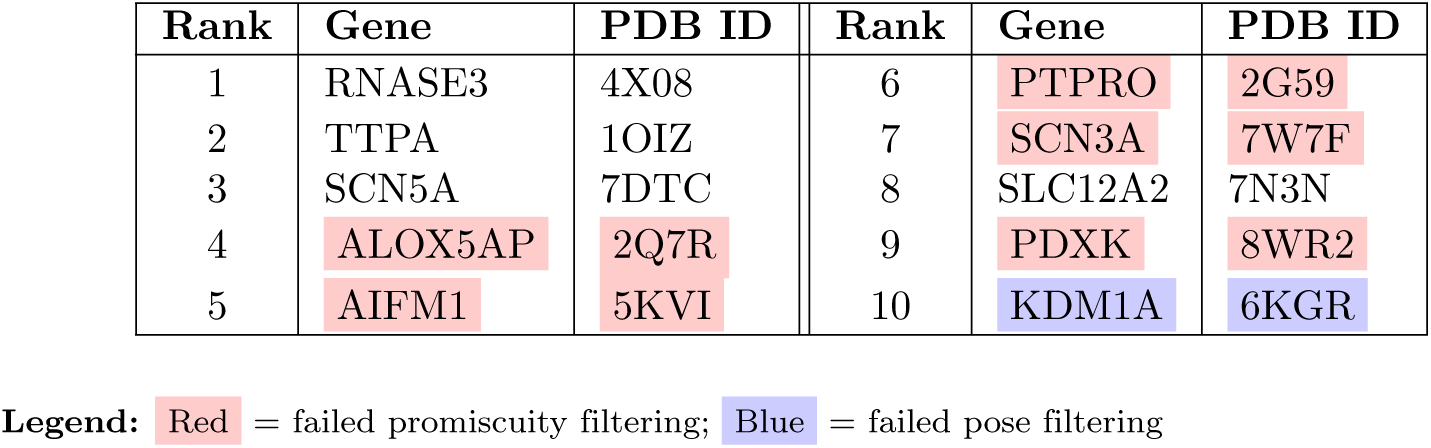
Paclitaxel off-targets identified after applying 25% promiscuity filtering (all 20 ligands) and pocket analysis validation. The known target TUBB ranked poorly at position 5724. Red highlighting indicates broad-binding targets removed by promiscuity filtering, blue highlighting indicates failed pose filtering.

| Rank | Gene | PDB ID | Rank | Gene | PDB ID |
| --- | --- | --- | --- | --- | --- |
| 1 | RNASE3 | 4X08 | 6 | PTPRO | 2G59 |
| 2 | TTPA | 1OIZ | 7 | SCN3A | 7W7F |
| 3 | SCN5A | 7DTC | 8 | SLC12A2 | 7N3N |
| 4 | ALOX5AP | 2Q7R | 9 | PDXK | 8WR2 |
| 5 | AIFM1 | 5KVI | 10 | KDM1A | 6KGR |

**Legend:** Red = failed promiscuity filtering; Blue = failed pose filtering

**1. Validated On-Target:** The intended target, BRAF V600E (Rank 6) was robustly identified by the ProteomeScan framework following promiscuity and pose filtering. The mutant form exhibited a much higher predicted binding affinity relative to wild-type BRAF (rank 90), with 93.8% pocket occupancy, indicating strong druggable site engagement. These computational findings are consistent with experimental data demonstrating selective inhibition of the MEK/ERK signaling pathway and cytotoxicity in BRAF V600E-positive cells [77]. This confirms accurate identification of dabrafenib’s primary therapeutic target.

**2. Literature Supported Off-Targets:**

- CYP1A2 (Rank 3): A hydroxy-dabrafenib metabolite (with a hydroxyl group attached to dabrafenib) is known to inhibit CYP1A2 (IC_50_ = 83 *µ*M) [59].This known related binding interaction supports our predicted interaction and suggests potential metabolic modulation by dabrafenib or its derivatives.

**3. Computationally Predicted Targets without Experimental Validation:**

- FABP2 (Rank 1): FABP2 is an intestinal fatty acid binding protein with a large hydrophobic cavity. While its primary function is dietary lipid absorption, FABP2 can bind diverse lipophilic molecules including at least one drug (the NSAID ketorolac) [78]. Dabrafenib’s high lipophilicity (log P ∼5.2) makes interaction with FABP2’s hydrophobic cavity plausible. Although no direct experimental evidence is available, the strong predicted binding affinity combined with FABP2’s affinity for lipophilic ligands suggests this interaction may be plausible.
- NF1 (Rank 10): NF1 is a known modulator of MAPK signaling and an established mechanism of dabrafenib resistance [79]. No direct dabrafenib interaction data exists.
- PROCR, CD1A, ACAD8, and SLC6A4 (Ranks 2, 7, 8, 9): While these proteins showed favorable docking characteristics, no experimental data currently support direct dabrafenib binding.

##### 6.3.2 Case Study 2: Trametinib

Similar to previous case study analysis, we started with a 25%(20) promiscuity threshold. This threshold ranked the MAP2K2/MAP2K1 targets at positions 361 and 521 respectively out of 7491 targets even after applying the promiscuity filter. We also experimented with running ProteomeScan under a stricter 10%(5) promiscuity threshold and pose filtering discussed in appendix. Under these conditions, known MAP2K2/MAP2K1 targets for trametinib were ranked at positions 60 and 130, respectively, out of 7,014 targets after promiscuity filters.

###### Target Analysis

This section summarizes the analyses of predicted targets identified for trametinib using the ProteomeScan framework.

**1. Validated On-Target:**

- MAP2K1/2 (Rank 521, 361): Trametinib binds directly to MAP2K1/2 with high potency, blocking MEK activation and downstream signaling. [80, 81]. The relatively poor ranks of these complexes were because of computational limitations involving complex interactions discussed in *Computational Limitations* section below.

**2. Computationally Predicted Targets without Experimental Validation:**

- PARP2 (Rank 2): MAP2K inhibition by trametinib downregulates BRCA2 protein expression, sensitizing cells to PARP inhibitors [82]. While PARP inhibitors such as olaparib block PARP2 to induce cell death [83], our computationally strong trametinib-PARP2 binding prediction suggests an alternative mechanism producing a similar phenotypic effect through direct PARP2 inhibition.
- BRAF V600E (Rank 3): BRAF V600E phosphorylates MAP2K1/2 [84]. The V600E mutation modulates inhibitor binding affinity but does not alter binding site accessibility [85]; thus, compounds that bind wild-type BRAF likely also bind BRAF V600E. Trametinib showed no direct BRAF binding in kinase selectivity screens [80], but it engages BRAF interface residues within BRAF/MAP2K ternary complexes [86]. Our computational prediction likely reflects this ternary complex mechanism.
- FABP2, CASP1 (Ranks 1, 4): While these proteins showed favorable docking characteristics, no experimental data currently support direct trametinib binding.

###### Computational Limitations

The ranking of known targets (MAP2K1/2) reflects fundamental computational challenges. Recent structural studies reveal that trametinib’s binding mechanism involves complex interactions with KSR proteins, where functional binding requires KSR1:MAP2K1 and KSR2:MAP2K1 complexes rather than isolated MAP2K proteins [87]. Traditional docking algorithms using static, isolated protein structures cannot model these assembly-dependent binding requirements.

Furthermore, trametinib demonstrates exceptional potency (IC50 0.7-0.9 nM) through allosteric mechanisms involving induced conformational changes that static docking approaches cannot capture. The disconnect between experimental potency and computational rankings validates the known limitations of structure-based approaches for conformationally complex targets.

##### 6.3.3 Case Study 3: Paclitaxel-TUBB

This case study highlights limitations of ProteomeScan and structure-based docking approaches in general when applied to assembly-dependent binding mechanisms, as the known target TUBB ranked poorly at position 5724 of 7657 targets. The failure stems from paclitaxel’s requirement for binding to assembled microtubules rather than isolated tubulin dimers, demonstrating that static docking methods fail to capture the cooperative conformational changes and assembly-state-dependent binding sites essential for this interaction.

###### Target Analysis

This section analyzes predicted targets identified for paclitaxel using ProteomeScan.

Computational results were compared against experimental and clinical literature to evaluate the biological relevance of predicted interactions following promiscuity filtering and pocket analysis.

**1. Validated On-Target**

ProteomeScan demonstrated poor predictive performance for the paclitaxel-TUBB interaction, yielding docking scores of -5.485 for 8V2J and -7.471 for 8V2J multi-chains. Most significantly, the known target TUBB ranked at position 5724 based on docking score, representing a failure in target identification. This result is notable given paclitaxel’s validated clinical efficacy as an anticancer agent and highlights fundamental limitations inherent in current structure-based computational approaches which have been discussed in *Mechanistic Basis for Prediction Failure* section below.

**2. Computationally Predicted Targets without Experimental Validation:**

1. SCN5A (Rank 3): Paclitaxel stabilizes microtubules to arrest cell division. SCN5A/NaV1.5 is a voltage-gated sodium channel normally expressed in cardiac tissue but aberrantly upregulated in triple-negative breast cancer, where it drives tumor cell proliferation. Blocking SCN5A improves paclitaxel’s effectiveness in TNBC models [88]. Paclitaxel is known to affect sodium channel family members indirectly through microtubule-mediated alterations in vesicular trafficking [89]. There is no evidence of a direct Paclitaxel-SCN5A interaction and this would require computational validation.
2. RNASE, TTPA and SLC12A2 (Ranks 1, 2, 8): These targets lack documented paclitaxel interactions and represent computational hypotheses requiring targeted validation.

###### Mechanistic Basis for Prediction Failure

The primary reason for the computational prediction failure resides in the assembly-dependent binding complexity of paclitaxel to tubulin. The taxane binding site in unassembled tubulin dimers is occluded by *β*M-loop dynamics, so the binding site is not fully formed or accessible in isolated tubulin units [36]. There is a significant 100-1000 fold affinity difference between paclitaxel binding to assembled microtubules (*K_D_* ∼10-50 nM) [37] versus free tubulin dimers (*K_D_* ∼1-10 *µ*M) [38], indicating that the assembly state is crucial for high-affinity binding. Furthermore, cooperative conformational changes across multiple structural elements of tubulin are required for paclitaxel binding, a complex process that static docking methods struggle to model accurately.

###### Computational Limitations

Molecular docking algorithms, including AutoDock Vina used in ProteomeScan, typically operate under a rigid receptor assumption, which cannot model the large-scale conformational reorganization necessary for paclitaxel binding. These methods rely on static structures, inherently missing the formation of the assembly-dependent binding site. Additionally, the scoring functions employed by these algorithms often poorly represent the complex cooperative binding energetics involved in systems like paclitaxel-tubulin interactions. This case study reveals the expected boundaries of structure-based approaches for targets that necessitate dynamic conformational transitions, assembly-dependent binding sites, complex allosteric mechanisms, and cooperative binding involving multiple protein states. This limitation is well-documented in the field of computational drug discovery [40], emphasizing that for such conformationally dynamic systems, experimental validation remains essential.

#### 6.4 Alphafold-based Analysis

We performed gene guided protein-ligand docking using the first 5000 targets from the full human proteome gene list. A set of five ligands (drugs) was selected for this analysis: Trametinib, Binimetinib, Dabrafenib, Vemurafenib, and TAK-632. Docking simulations were conducted using AutoDock Vina with 32 exhaustiveness and mode 8 settings. For target structures, AlphaFold-predicted PDB files were used. It’s important to note that AlphaFold provides a pLDDT (predicted Local Distance Difference Test) score for each residue, which ranges from 0 to 100. Higher pLDDT values indicate greater confidence in the structural prediction and typically better model accuracy.

Focusing on Trametinib as an example, docking attempts were made across the AlphaFold-predicted structures of 5000 genes. Out of these, approximately 2000 docking runs were successful. Among the successful cases, only 1276 genes had corresponding experimental PDB structures available for comparison. Most of these experimental PDBs had sequence lengths in the range of 300 to 400 residues, and docking scores between the experimental and AlphaFold structures showed only minor differences within this range.

However, a key limitation observed was the failure of docking runs for AlphaFold PDBs with sequence lengths exceeding 500 residues. The success rate of docking correlated with two primary factors: shorter sequence lengths (generally below 500 residues) and higher average pLDDT scores. In contrast, failed docking runs were often associated with larger sequence lengths and lower pLDDT scores, indicating lower confidence in the predicted structures. An example is the gene EXPH5 (UniProt ID: Q8NEV8), which has a sequence length of 1989 residues and a low per-residue confidence score for a large section of the structure, leading to docking failure with limited compute.

Overall, AlphaFold PDBs with sequence lengths in the 300–500 range tend to yield successful docking results under the current AutoDock Vina setup, with minimal deviation from docking scores obtained using experimental PDBs. In contrast, large AlphaFold structures not only have lower average pLDDT scores but also show higher failure rates in docking, making their predicted top docking hits potentially unreliable. Additionally, docking scores from experimental PDBs generally outperform those from AlphaFold structures, particularly when comparing targets with smaller sequence lengths.

#### 6.5 Promiscuous Targets

Thresholding the top 25% targets common in all 20 ligands gives the following 166 promiscuous targets: DYRK2, DGAT1, CAMK2A, FN3K, ASAP3, LIN7C, LARGE1, KIF5A, DHFR, AMPD2, CCR5, NR5A2, SLC7A11, CARM1, ARAP3, BCHE, PTPRO, IDO1, PHF8, PRKCQ, HTR2A, GLUL, ALDH1L1, ALOX5AP, ADH5, SNRNP70, DOT1L, CHRM1, MICU1, VKORC1, CYP3A5, KDM1A, AKR1B1, PTGR2, OAT, HNMT, PIKFYVE, MMP12, F11, ADH4, GSTK1, CFTR, BHMT, FECH, ALDH1A2, ERAP1, NUMB, GPR156, CBX1, PDE4D, ALDH3A1, FGFR2, IZUMO1R, MFSD10, ATG9A, AIM2, NUP155, LNPEP, TRIAP1, SIRT6, PTK2B, CYP3A4, FAM111A, FLNA, NPC1, PDE4B, HSP90AA1, AIFM1, KLKB1, DCPS, SOAT1, GHSR, HCRTR1, DPP3, AOX1, IAPP, HSD17B4, NQO1, NMT1, PDE4A, AGTR2, AFM, HTR2B, SETD3, CYP21A2, MAT1A, BST1, PML, FARS2, ADORA2A, DMGDH, PDXK, PARP3, LDHA, PYCARD, AGER, SCN3A, MAML1, MGST2, MAT2A, FOLH1, ACLY, KYAT1, HDAC8, HHIP, GBA3, AQP10, IRF3, EPC1, GPR155, GM2A, SETDB1, CASP1, GLS, CDKL1, KDM5A, NTAN1, IQGAP1, ENPP2, PADI3, FGL1, CSNK1E, SQLE, KMO, SAT2, NOS1, CD1E, GABARAP, CPTP, CETP, SOAT2, ALDH1A3, BPHL, SRPK1, SLC14A1, CASQ1, AKR1B10, FZD2, ACAD9, GMDS, CSNK2A1, MFSD2A, RIPK2, HHAT, ACHE, HCRTR2, RTN4IP1, SLC2A3, ATXN3, ADCY10, ACADM, AKR1C3, ADAM22, PCIF1, NTRK3, ALDH3A2, SLC9A3, CACNA1E, MELK, CHIA, BACH1, CARNMT1, PROCR, GSTA1, TUT7, AOC1.

#### 6.6 Pseudo-code

This section contains details of algorithms used for ProteomeScan:

- Algorithm 1: Proteome Scan algorithm
- Algorithm 2: Get optimal cleaned PDBs per Gene
- Algorithm 3: Select optimal PDBs
- Algorithm 4: Clean the PDB file
- Algorithm 5: Run gene guided protein-ligand docking
- Algorithm 6 Analyze target promiscuity
- Algorithm 7 Check binding (pose analysis)

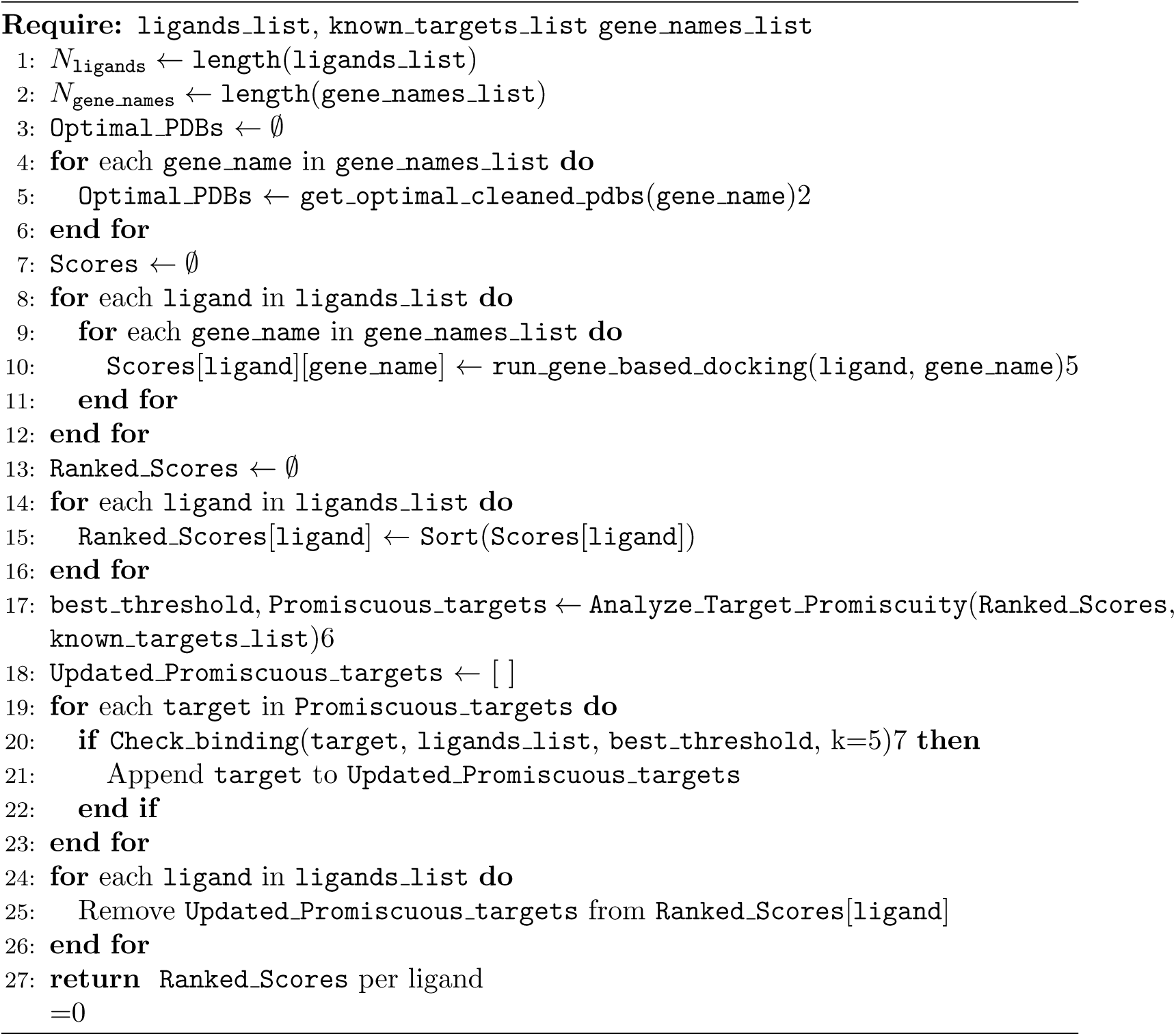

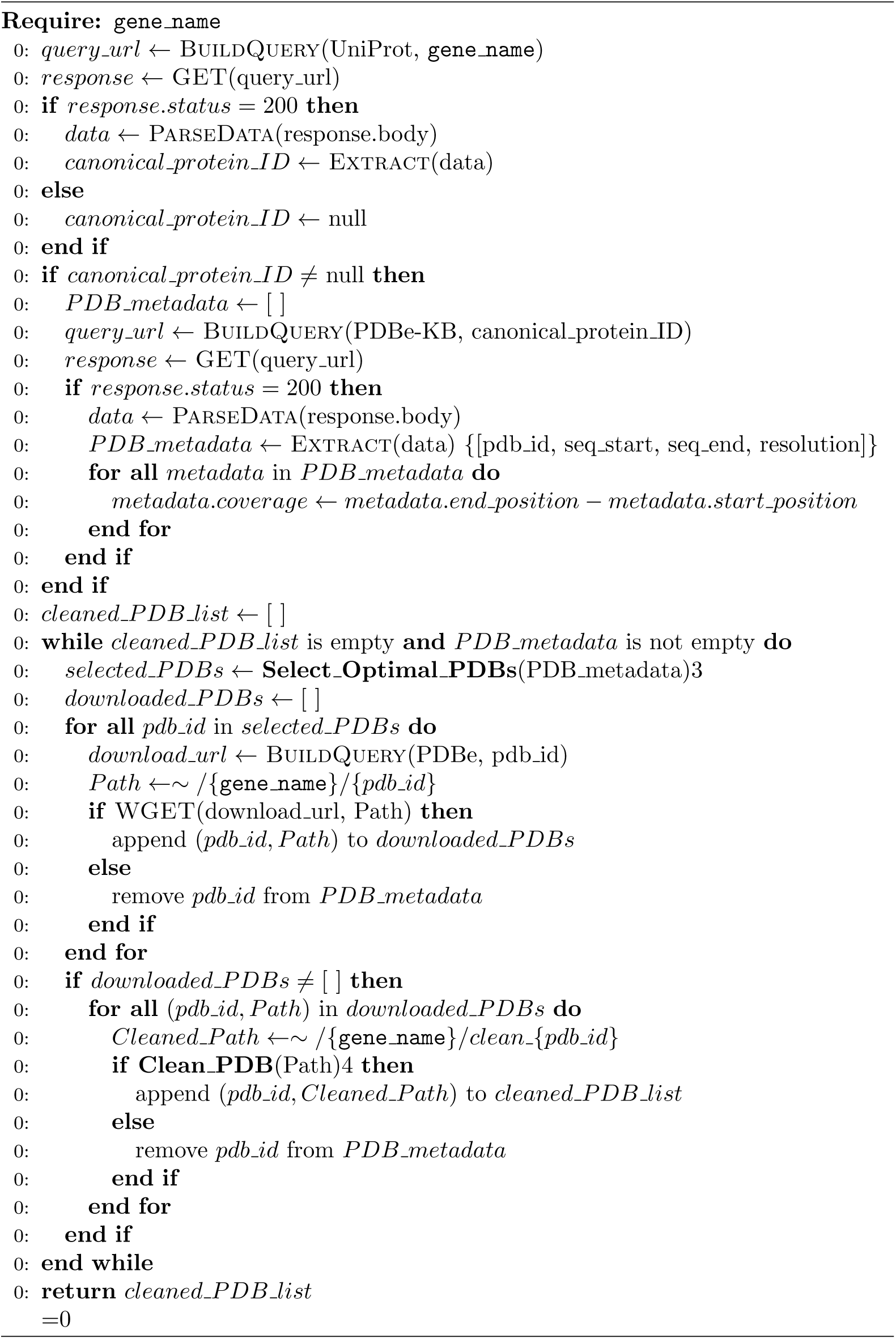

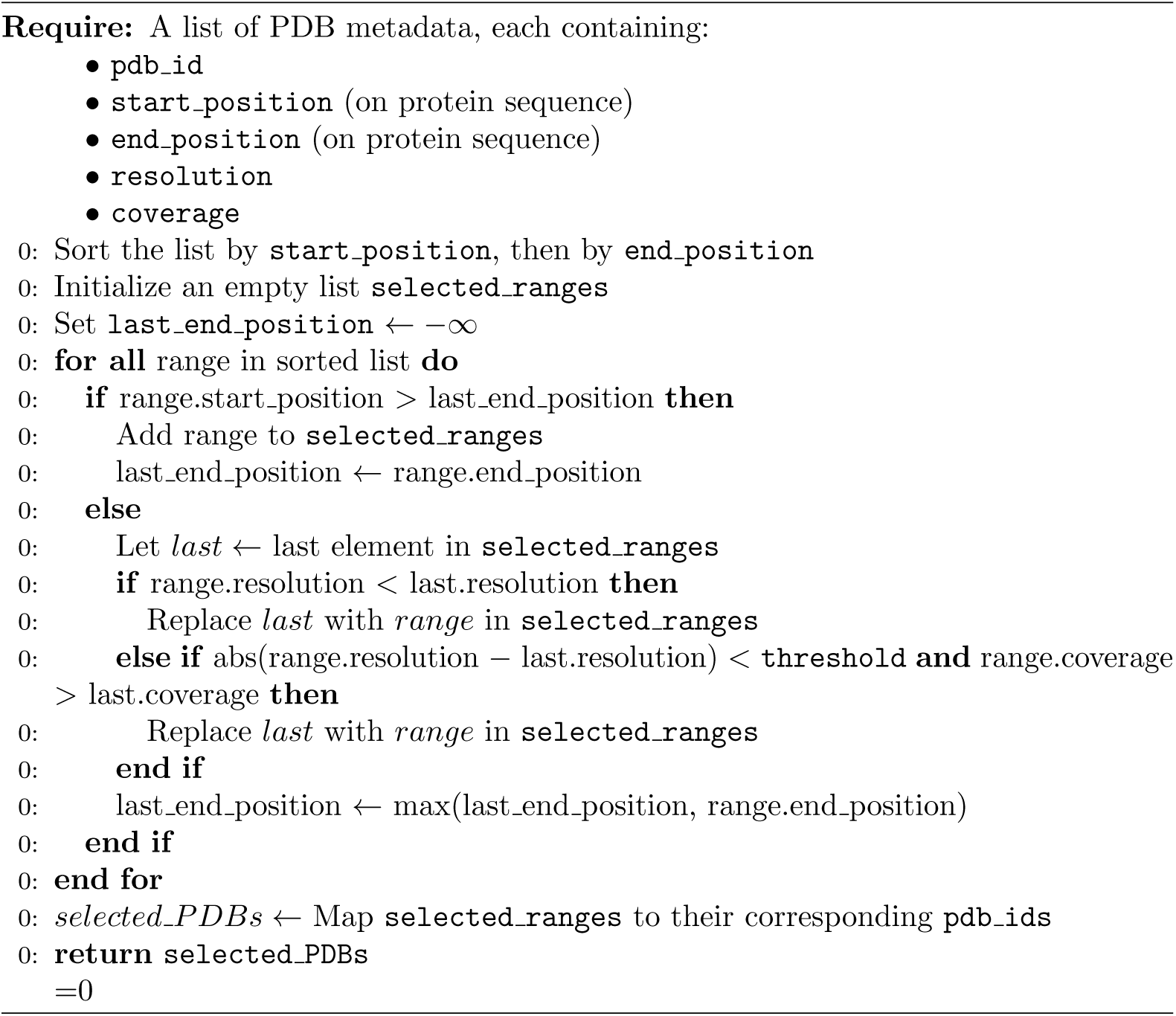

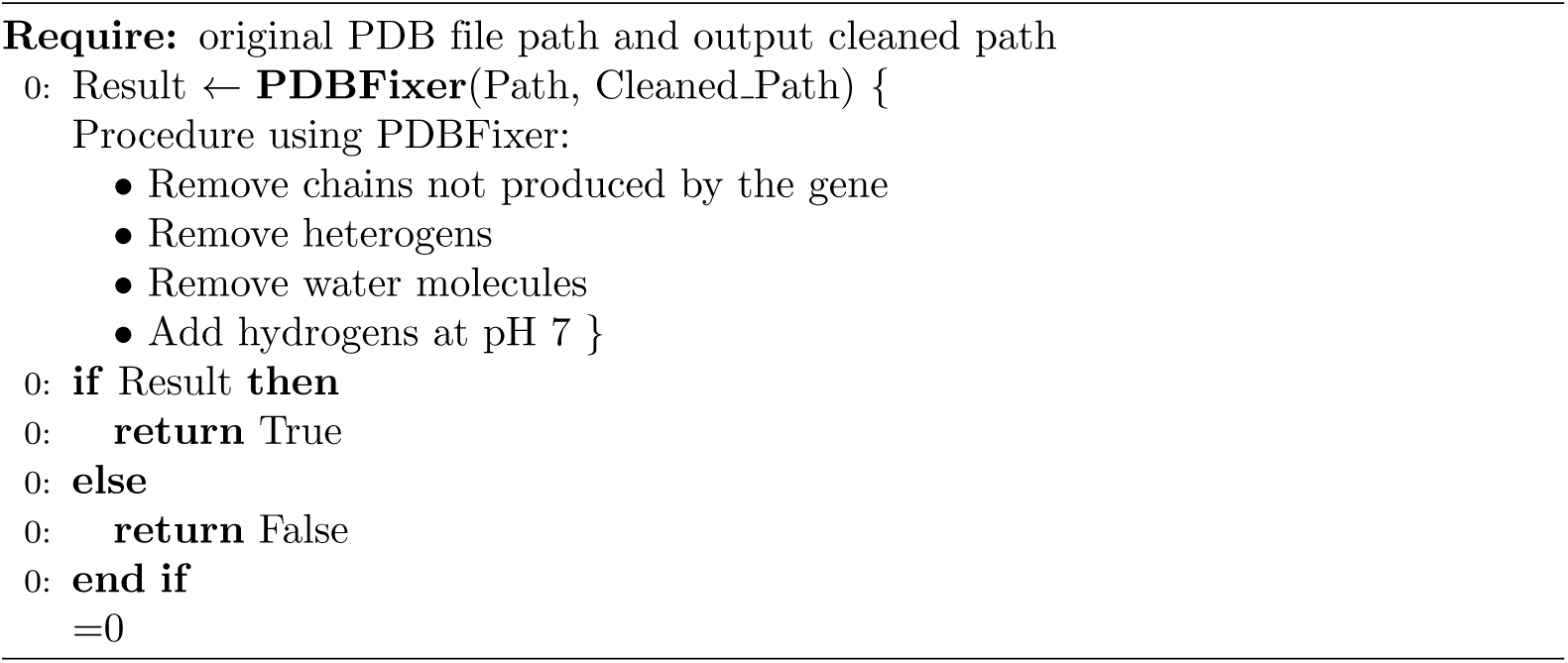

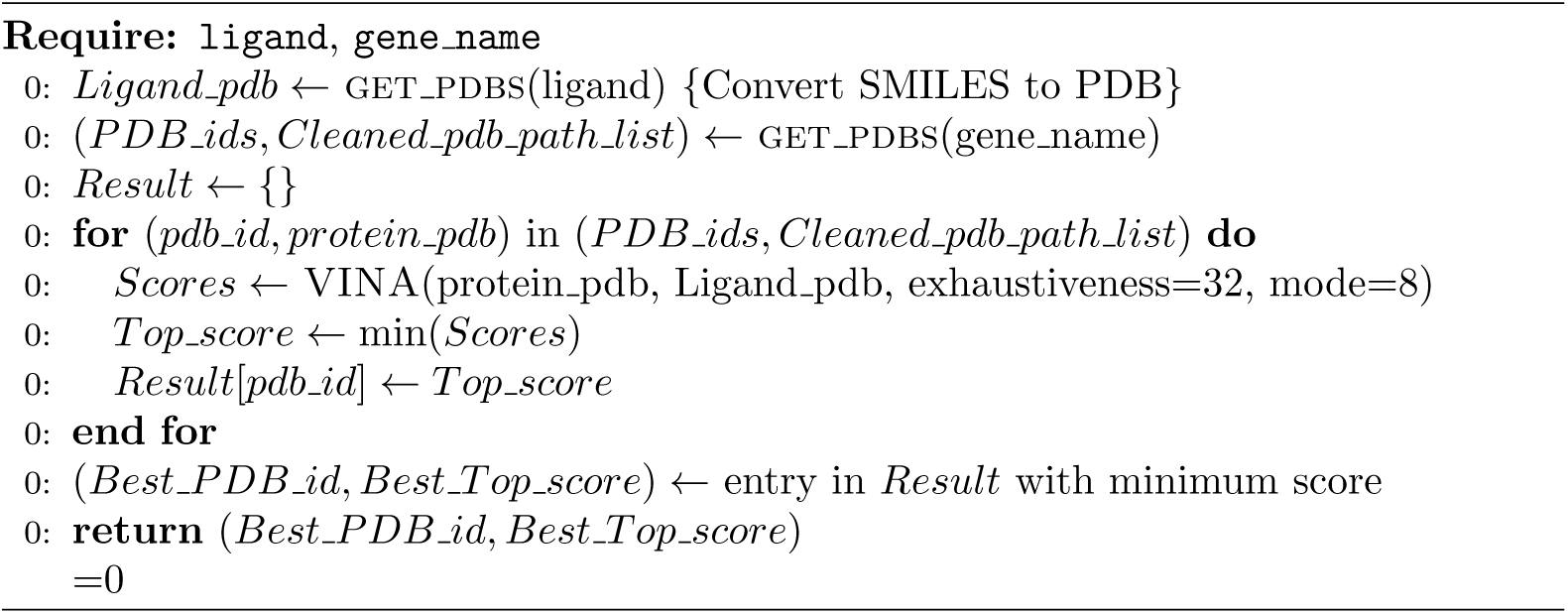

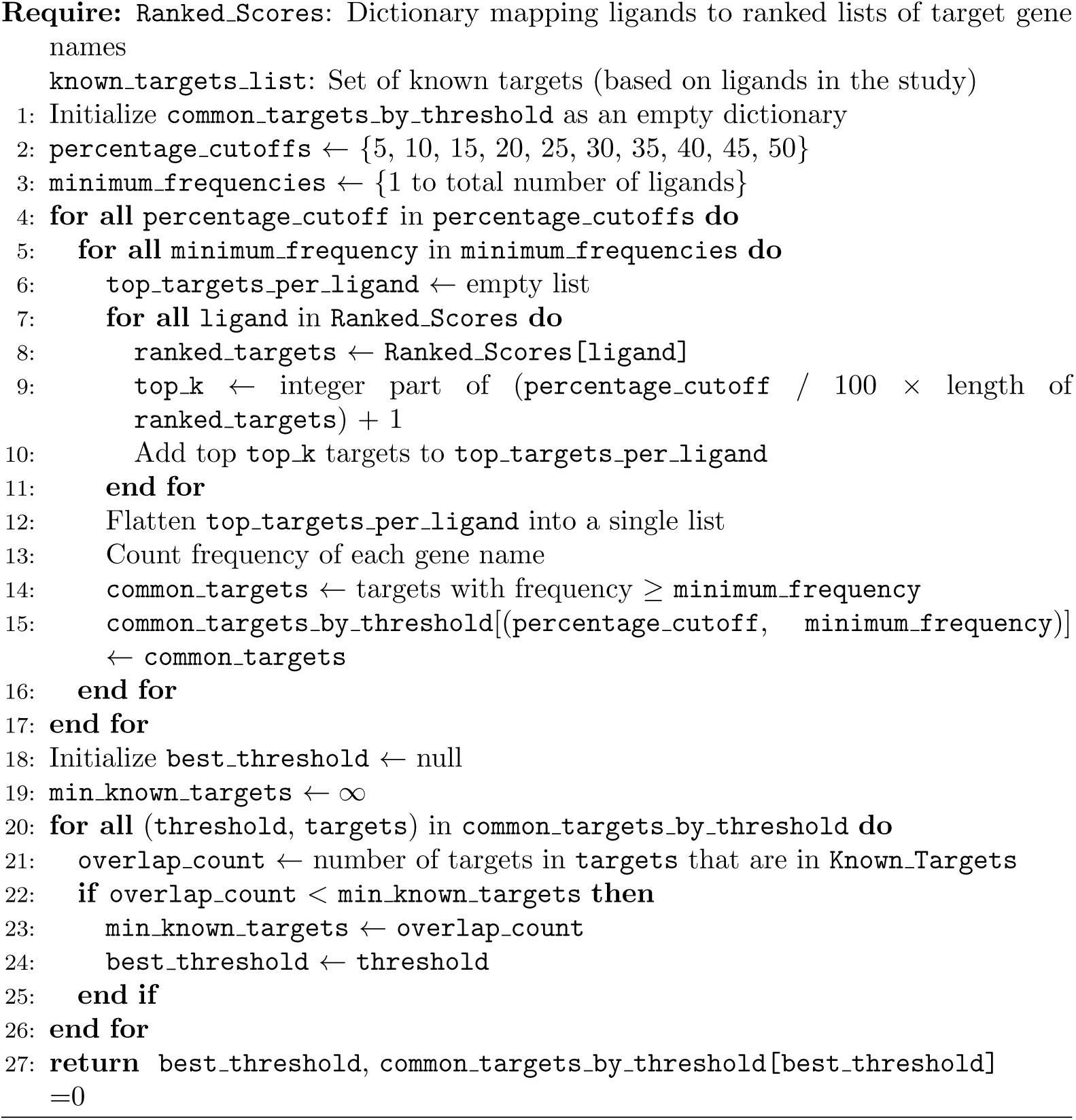

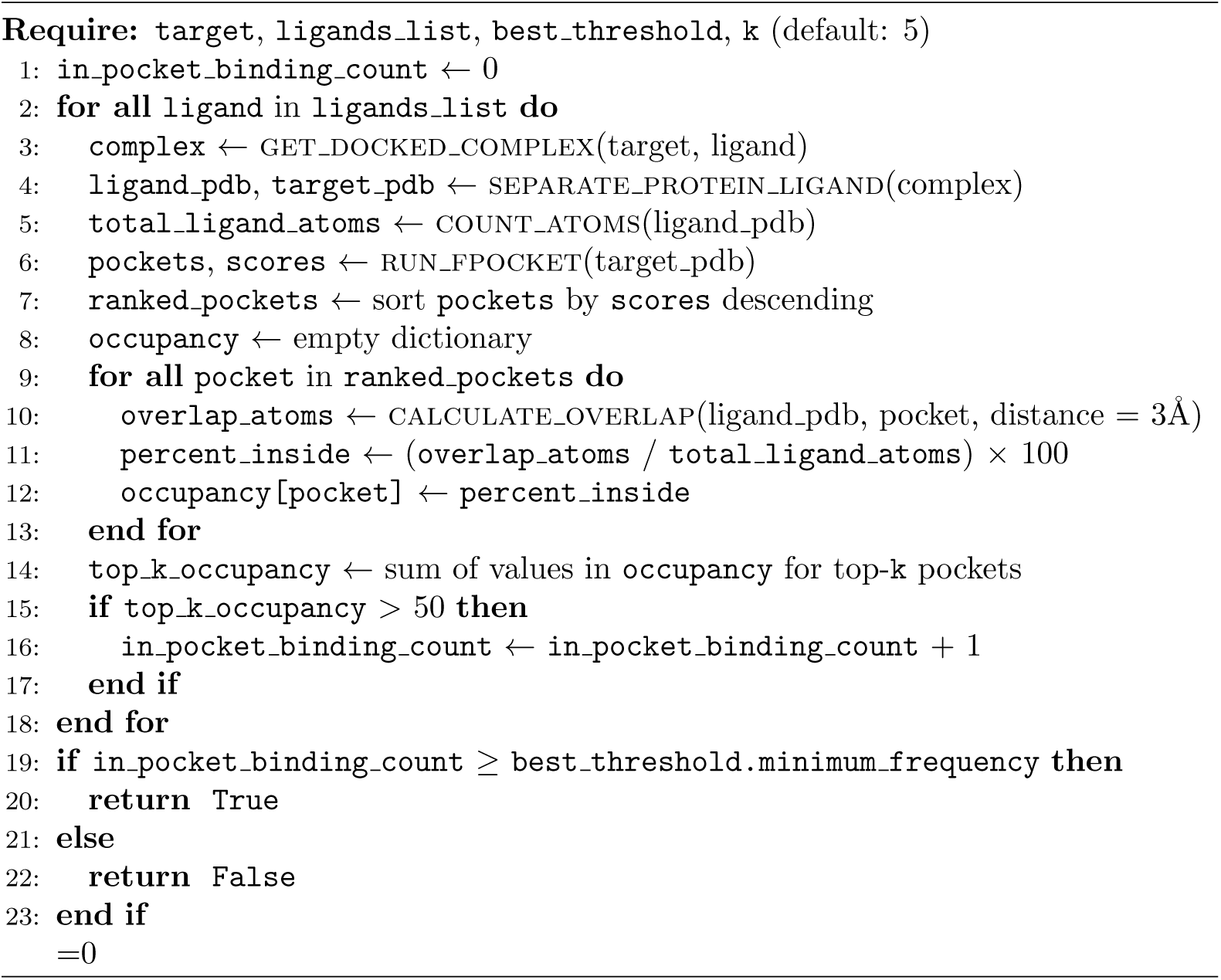

